# Staufen2 modulates the temporal dynamics of human neurogenesis *in vitro*

**DOI:** 10.1101/2025.10.02.679988

**Authors:** Akshay Jaya Ganesh, Sandra María Fernández-Moya, Ana Gutiérrez-Franco, Natalie C. Ferreira, Rafael Tur-Guasch, Damià Romero-Moya, Loris Mularoni, Alessandra Giorgetti, Monika Piwecka, Agnieszka Rybak-Wolf, Mireya Plass

## Abstract

RNA-binding proteins (RBPs) play a central role in post-transcriptional regulation during brain development, yet their specific functions in coordinating human neural lineage decisions remain poorly understood. Here, we investigate for the first time the role of the double-stranded RBP Staufen2 (STAU2) in human neurogenesis. Characterization of STAU2 knockout iPSC derived cells using scRNA-seq shows that loss of STAU2 disrupts neuroepithelial cell identity and accelerates neural differentiation by altering the activity of key transcription factors and driving early metabolic transitions. Additionally, STAU2 regulates the expression of miRNA host genes and alters miRNA-mediated post- transcriptional control in progenitor cells, which exerts additional effects on STAU2 regulated gene regulatory networks. These changes result in neural progenitor exhaustion, unstructured neural rosettes, and reduced organoid size. Together, our work uncovers a previously unrecognized role for STAU2 as a central regulator of early human neurogenesis, acting through both miRNA-mediated and transcriptional pathways to coordinate progenitor maintenance and neuronal fate specification.

## Introduction

Neurogenesis is crucial for brain development and function during embryogenesis^1^ and throughout life^2,3^. This process involves neuroepithelial cells —the earliest neural stem cells (NSCs)— which differentiate into specialized cell types that integrate into neural circuits^1^. While the genetic regulation of neurogenesis by pro-neural transcription factors has been extensively studied^4,5^, recent work highlight the importance of post-transcriptional regulation in fine-tuning neurogenesis and shaping cortical development. RNA-binding proteins (RBPs) interact with target RNAs to regulate key post-transcriptional events, including splicing, RNA stability, localization and translation^6–12^. Dysregulation of RBPs during brain development impairs neuronal migration and synaptic plasticity, leading to behavioral alterations and contributing to both neurodevelopmental and neurodegenerative disorders^13–15^. The Staufen (Stau) family of double-stranded RBPs is essential for neurogenesis and neuronal function across species^7,16,17^. Initially described in *Drosophila,* Staufen mutants disrupt *prospero* mRNA localization in mitotic neuroblasts, impairing proper cell fate specification^18^. Mammals express two Stau orthologs: *Staufen 1* (*Stau1*), which is ubiquitously expressed, and *Staufen 2* (*Stau2*), which is enriched in the brain^19^. During mouse brain development, cortical radial glial cells (RGCs) undergo repeated asymmetric cell divisions to produce both self-renewing progenitors and committed neural progenitor cells^20^. STAU2, together with two other RBPs, Pumilio 2 (PUM2), and DEAD-box helicase 1 (DDX1), is asymmetrically distributed in RGCs, enriched in TBR2+ daughter neuroblasts together with specific neuronal RNAs^16,17^. Loss of STAU2, PUM2 or DDX1 in the developing mouse cortex triggers premature RGC differentiation and mislocalization of key mRNAs such as *Prox1* and *Trim32*^16^, suggesting a role in intermediate progenitor cell (IPC) fate determination via asymmetric RNA distribution. Interestingly, STAU2 protein is expressed even earlier in mouse neurogenesis^21^ during symmetric neuroepithelial cell proliferative divisions to expand the neural stem cell pool^22^.

Beyond mRNA localization, STAU2 has also been implicated in Staufen-mediated mRNA decay (SMD), an mRNA degradation process that is mediated by the binding of Staufen proteins to a STAU-binding site (SBS) within the 3’-untranslated region (3’UTR) of target mRNAs^23^. Despite these insights, which have been derived from animal models, the function of STAU2 in early human neurogenesis, particularly in regulating distinct sets of mRNAs during differentiation from neuroepithelial to neural stem cells, remains unknown.

Recent advances in stem cell technologies have enabled study neurogenesis *in vitro* using induced pluripotent stem cells (iPSCs)^24^. iPSCs can differentiate into a wide variety of cortical neuron subtypes *in vitro*^25,26^, recapitulating critical stages of human corticogenesis — such as neuroepithelial formation, RGC specification, and neuronal maturation^27^. Thus, they provide a robust model to study early neurogenesis and to elucidate the molecular mechanisms that govern neural fate specification and maturation in humans.

In this study, we employed single-cell transcriptomics to investigate the role of STAU2 for the first time in a human context at four key stages during neurogenesis *in vitro*: the formation of the neural epithelia, neural progenitor cell (NPC) proliferation, neural specification, and neuronal maturation. Our results show that STAU2 knockout (KO) induces transcriptional and metabolic reprogramming during early human neurogenesis, with particularly strong effects in neuroepithelial cells. We observed a coordinated downregulation of translation and RNA-processing pathways, coupled with an enrichment of glycolysis, oxidative phosphorylation, and cholesterol/steroid biosynthesis programs typically associated with neuronal differentiation. This shift destabilizes progenitor identity, leading to premature neuronal commitment and early depletion of the progenitor pool.

Beyond metabolism, STAU2 KO cells displayed enrichment of gene sets related to neurotransmitter receptor signaling, synaptic function, and axon guidance across multiple neuronal clusters, suggesting accelerated acquisition of neuronal functional signatures. At the regulatory level, we identified reduced activity of transcriptional regulators ARID3A and CHD2, which normally sustain chromatin accessibility and maintain progenitor states, together with altered expression of multiple miRNA host gene. This included miR-9 and miR- 218, key regulators of cortical development, implicating STAU2 in the integration of transcriptional and miRNA mediated network.

Together, our findings underscore the pivotal role of STAU2 as a central post-transcriptional hub that coordinates gene expression, metabolism, and regulatory network activity to align progenitor maintenance with the temporal dynamics of neurogenesis, thereby shaping the architecture of the developing human brain.

## Results

### Generation of functional STAU2 knockout (KO) iPSCs using CRISP/Cas9

To investigate the role of STAU2 in human neurogenesis, we generated functional STAU2 KO iPSC lines using CRISPR/Cas9 genome editing. STAU2 is expressed as multiple alternatively spliced isoforms of approximately 52, 59, and 62 kDa, which differ in their N- and C-terminal regions^19^. To eliminate all functional proteins, we specifically deleted exons 7 and 8 of STAU2 gene, which partially encode the second and third double-stranded RNA binding domains (dsRBM2 & dsRBM3)^19,28^ **(Fig. 1A)**. This deletion disrupted the two dsRBM dimers, preventing STAU2 binding to the RNA, and introduced a premature stop codon, leading to reduced mRNA levels **(Fig. 1B)**. Consistently, the generated KO lines completely lacked the expression of canonical STAU2 protein isoforms **(Fig. 1C)**. Using this strategy, we successfully established two KO iPSC lines, STAU2-1 and STAU2-2, derived from two independent parental iPSC lines. We validated that both cell lines retained normal karyotypes **(Fig. S1)** and pluripotency, as demonstrated by their ability to differentiate into derivatives of all three germ layers **(Fig. 1D).**

**Figure 1.**
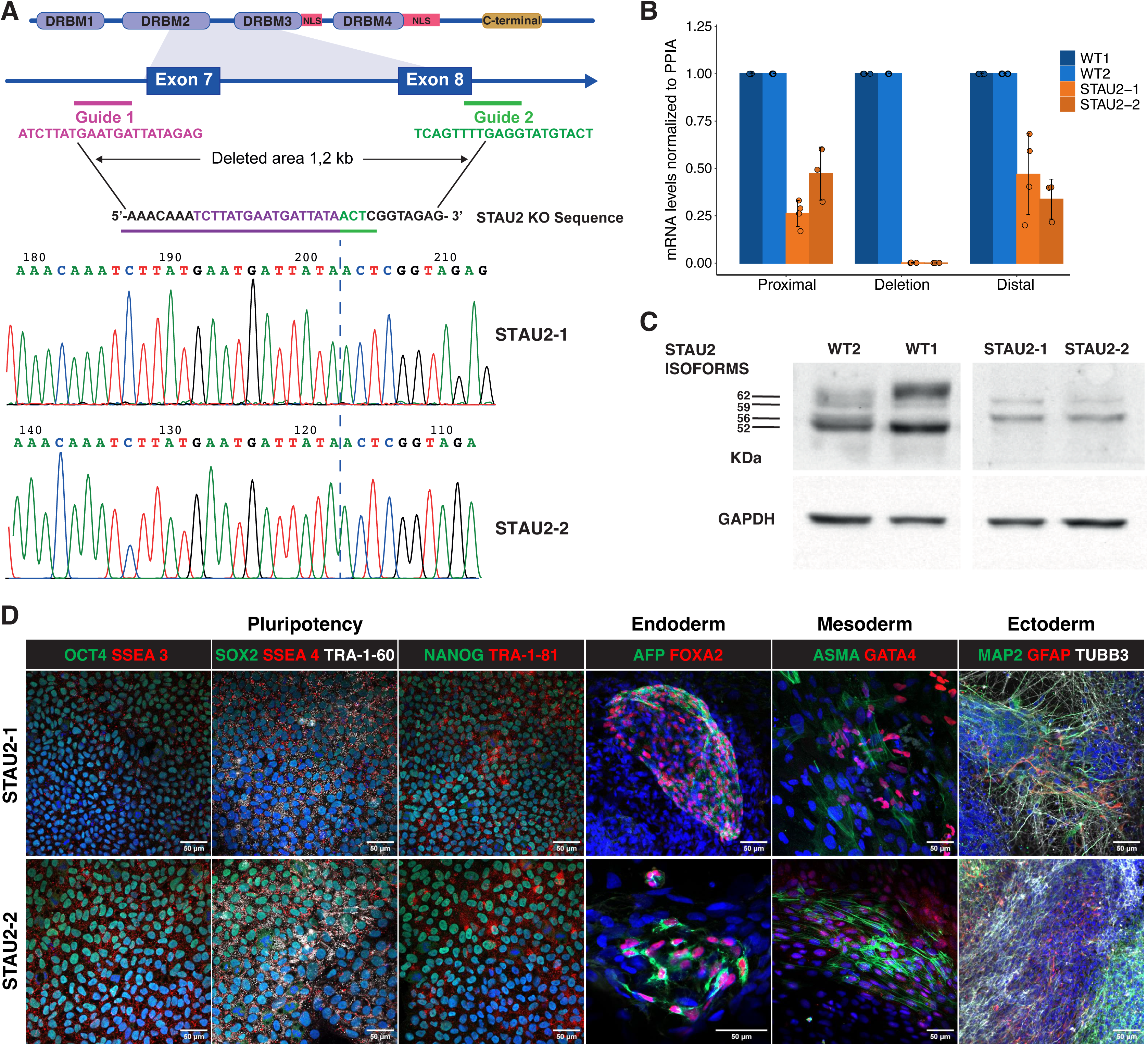
Generation and characterization of STAU2 knockout human pluripotent stem cells. **A.** Schematic of CRISPR/Cas9-mediated STAU2 knockout strategy targeting exons 7 and 8 partially deletes DRBM2 & DRBM3. Two guide RNAs flank a ∼1.2 kb deletion. Sanger sequencing confirms the deletion of STAU2-1 and STAU2-2 clones. **B.** RT-qPCR analysis of STAU2 mRNA levels using primers targeting regions proximal, within, and distal to the deleted area. Significant reduction is observed in the deletion and distal regions in knockout lines (n=3 independent replicates; mean ± SEM). **C**. Western blot analysis showing loss of STAU2 protein isoforms in both STAU2 knockout lines compared to WT control. The faint bands observed do not correspond to any known STAU2 isoforms, or the truncated protein predicted after deleting exon 7 and 8, and are likely due to nonspecific antibody binding. GAPDH serves as loading control. **D.** Immunofluorescence staining of pluripotency markers SSEA-4, SSEA-3, TRA-1-60, TRA-1-81, and the transcription factors OCT4, SOX2 and NANOG in both knockout clones. Differentiation into three germ layers is confirmed by expression of lineage-specific markers: AFP and FOXA2 (endoderm), ASMA and GATA4 (mesoderm), MAP2, GFAP, and TBB3 (ectoderm). Scale bars, 50 µm.

### STAU2 loss accelerates neuronal commitment and perturbs fate specification

We differentiated STAU2 KO cells from one of the generated cell lines (STAU2-1) and their isogenic control into neural and glial lineages using a protocol previously established in the lab^29^. Briefly, hiPSC colonies were induced toward a neuronal fate using a combination of BMP inhibitors (noggin, dorsomorphin, and SB431542). We collected samples at five timepoints (Day 0, 11, 25, 55 and 70) to characterize the cell populations obtained using single-cell transcriptomics **(Fig. 2A)**. Initial morphological characterization of the iPSC-derived populations along the differentiation did not reveal significant differences between STAU2 KO and WT lines. We detected proliferating neuroepithelial cells at day 11 (D11), characterized by the expression of SOX2 and PAX6. At day 25 (D25), we identified proliferating neural progenitor cells (NPCs), expressing PAX6 and KI67, and cortical progenitors expressing TBR1 (early-born deep-layer neurons) and BRN2 (upper-layer fate and intermediate progenitors). At this stage, we noticed a higher amount of TBR1+ cells in the KO than in the WT cell line, suggesting accelerated or premature neurogenesis. At day 55 (D55) and day 70 (D70), we detected MAP2+ mature neurons, TBR1+ and BRN2+ neurons, and astrocytes expressing GFAP **(Fig. 2B)**.

**Figure 2.**
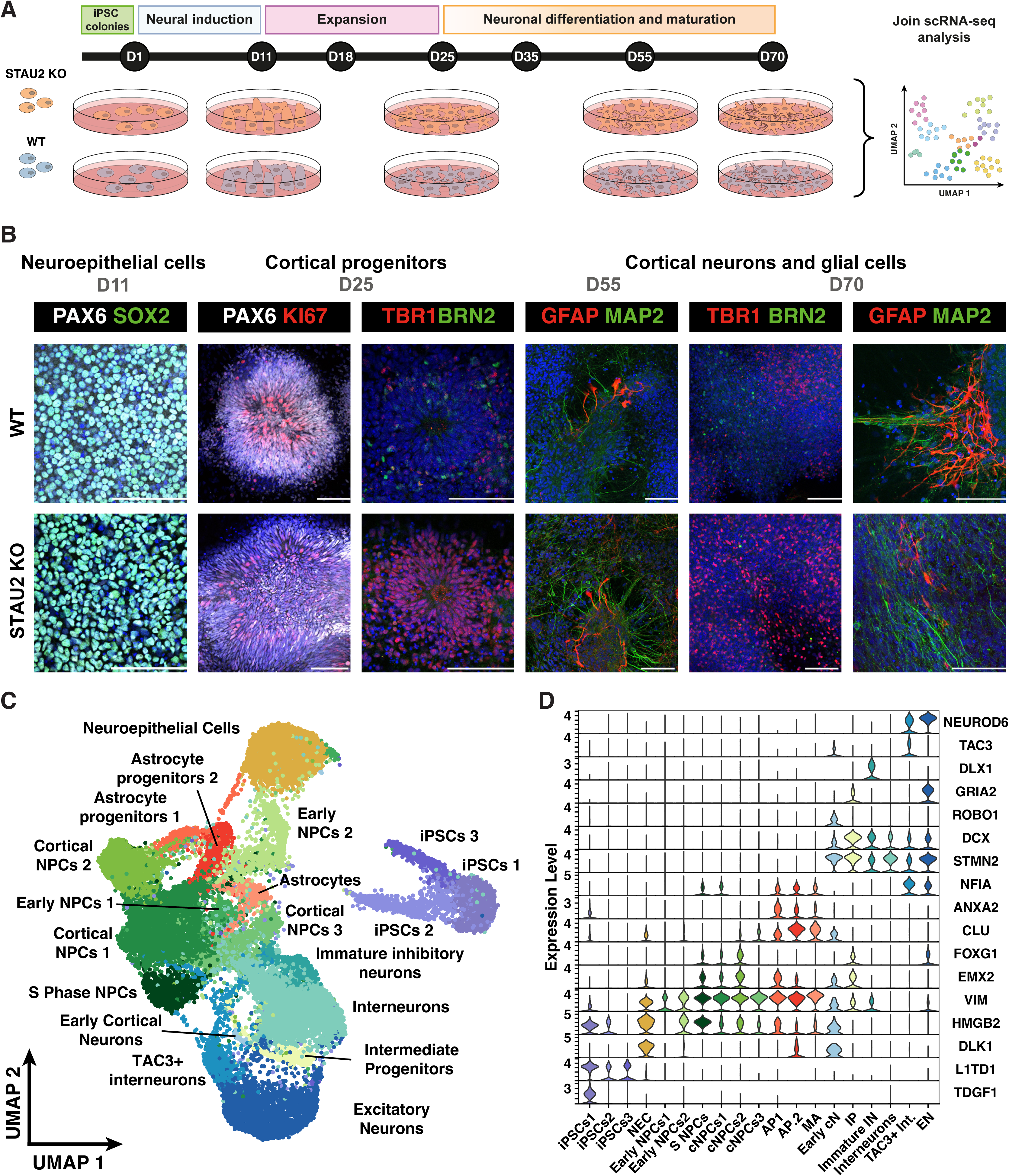
scRNA-seq to study effect of STAU2 KO throughout differentiation stages. **A.** Schematic representation of the differentiation of iPSCs and the experimental approach. **B.** Immunofluorescence validation of the neural differentiation at defined timepoints. Both WT and STAU2 KO lines form neuroepithelial progenitors (day 10, PAX6, SOX2), proliferating cortical progenitors (day 25, PAX6, Ki67, TBR1, BRN2), and cortical neuron/glial populations (from day 35 onward, TBR1, BRN2, MAP2, GFAP). STAU2 KO cells show earlier expression of neuronal markers. Scale bars, 100 µm. **C.** Annotated UMAP containing all the WT and STAU2 KO cells collected during the time-course differentiation. Distinct clusters represent key neurodevelopmental stages, including iPSCs, neuroepithelial cells, neural progenitor cells (NPCs), intermediate progenitors, differentiated astrocytes, interneurons and excitatory neurons. **D.** Violin plots showing the expression of known cell type markers across the different clusters including iPSC markers (TDGF1, L1TD1), neuroepithelial population (DLK1), progenitor populations (VIM) and astrocyte populations (CLU, ANXA2). Neuronal populations include early neurons (DCX), early cortical neurons (ROBO1), intermediate progenitors (EMX2), and excitatory neurons (NEUROD6, NFIA).

We performed single-cell RNA sequencing (scRNA-seq) on all these samples using Drop-seq. After quality control and filtering, we obtained a total of 29,368 cells and 27,138 genes, with an average of 1,162 genes and 2,158 UMIs per cell **(Fig. S2, Table S1)**. To compare all the data generated, we integrated the samples from all the time points from the different time-course experiments using Canonical Correlation Analysis (CCA) and performed joint clustering of all samples with Seurat v4^30^ **(Fig. S3)**. We used FindAllMarkers function from Seurat v5^31^ to identify top marker genes for each cluster **(Table S2, Fig. S4)**. Based on these markers, along with well-established lineage-specific markers **(Table S3)**, we annotated the cellular identities of each cluster. This analysis identified 19 clusters, representing various stages of neuronal differentiation including iPSCs, neuroepithelial cells, multiple NPC clusters, intermediate progenitors, interneurons and excitatory neurons, astrocyte precursors, and mature astrocytes **(Fig. 2C, D).** Most clusters aligned with the expected progression of neural differentiation, as observed in the macroscopic characterization of the cell cultures with imaging **(Fig. 2B, S5A)**. Interestingly, at D11, apart from neuroepithelial cells and NPCs, we identified a new cell population expressing markers characteristic of cortical neurons at a much earlier time point than anticipated. These early cortical neurons expressed markers such as NEUROD4 and NEFM **(Fig. S5B),** suggesting that at this early state some cells already are committed towards the neuronal fate. At D25, we identified committed progenitor populations, such as astrocyte progenitors, cortical NPCs, and intermediate progenitors, as well as early neurons and immature inhibitory neurons. By D55 and D70, further maturation was evident, with clusters corresponding to mature neurons, mature astrocytes, and residual progenitor populations **(Fig. S5A)**.

To assess fate specification and maturation quantitatively, we analyzed the cell composition of the scRNA-seq samples across time points using scCODA^32^. We found a significantly higher proportion of astrocyte progenitors at D25 and of excitatory neurons at D70 in STAU2 KO samples compared to control **(Fig. S6, Table S4)**. While these effects are significant and indicate shift in lineage specification, we did not observe a global increase in the abundance of mature cell types.

### Differential expression analysis reveals a strong impact of STAU2 on neuroepithelial cells

To assess the impact of STAU2 KO in individual cell populations, we performed differential expression analysis within each cluster using the MAST implementation in Seurat^31,33^. We chose MAST as it performs well for data with large batch effects, especially compared to pseudobulk approaches^34^. To address systematic variability in gene expression across batches, we used a mixed model to account for confounders such as batch, number of genes and UMIs, and the percentage of UMIs mapped to mitochondrial and ribosomal genes. Differential expression analysis revealed that the neuroepithelial cell cluster displayed by far the greatest number of significant differentially expressed genes (DEGs) between STAU2 KO and control **(Fig. 3A, Table S5).** As neuroepithelial cells represent the main cell population at D11 **(Fig. S5A)**, this suggest that the effect of STAU2 KO is stronger at the early progenitor stage. Other clusters including NPCs and neurons also showed gene expression changes between STAU2 KO and WT cells, but these were to a much smaller extent.

**Figure 3.**
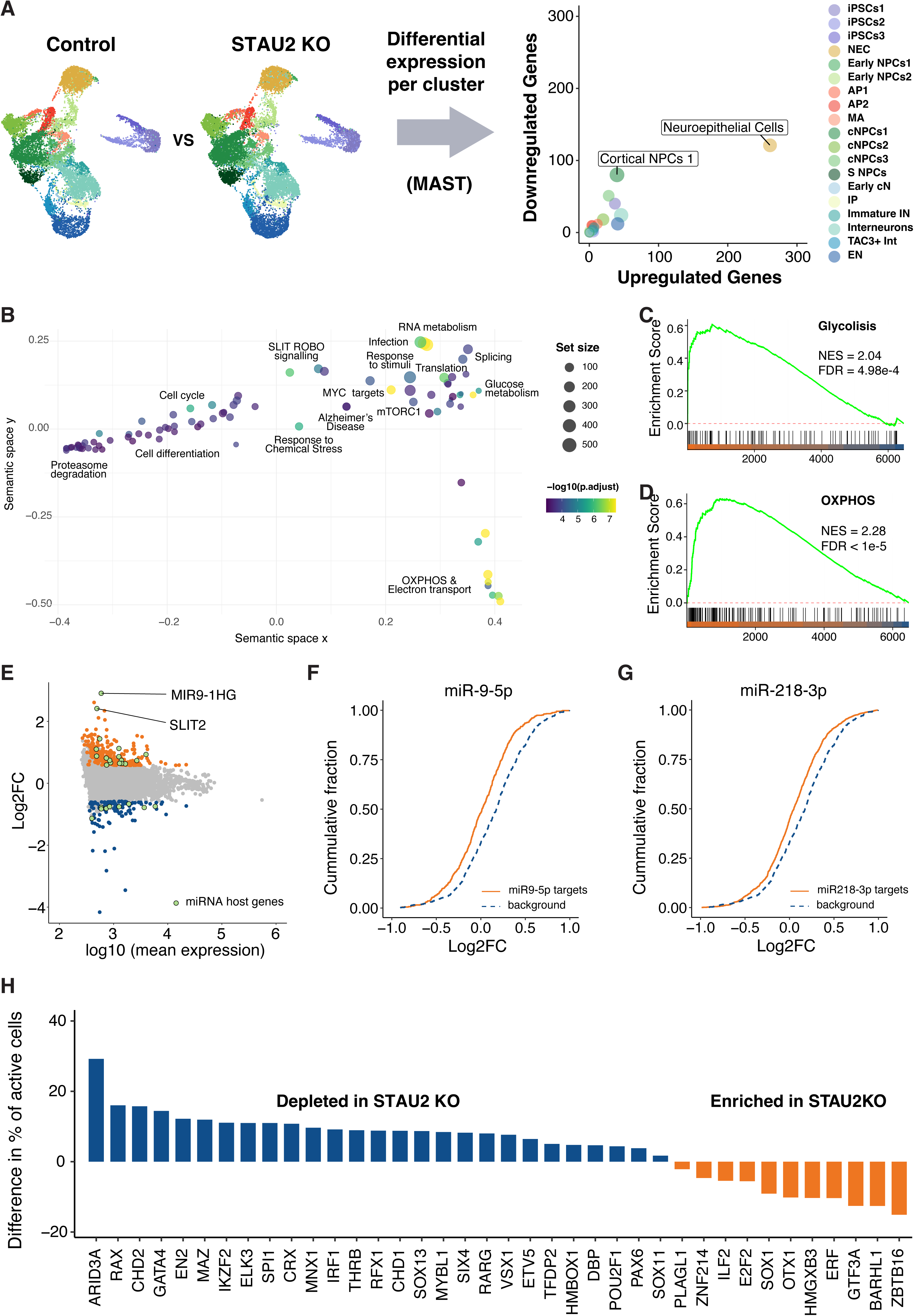
STAU2 KO affects gene expression through deregulation of miRNA host genes in neuroepithelial cells. **A.** Schematic representation of the cell-type specific differential expression analysis using MAST and scatterplot showing the number of differentially expressed genes (adjusted p < 0.05, |log₂FC| > 0.58, MAST test) in each cluster, comparing STAU2 KO to WT. Neuroepithelial cells and Cortical NPCs 1 show the highest number of transcriptional changes. Circle size indicates the number of cells per cluster**. B.** Gene set enrichment analysis (GSEA) of DEGs in STAU2 KO neuroepithelial cells shows significant upregulation of oxidative phosphorylation and glycolysis pathways (FDR < 1e-5 & FDR = 4.98e-4 respectively), as well as pathways related to DNA repair, hypoxia, mTORC1 signaling and ROS. These metabolic transitions are associated with cell cycle exit and neuronal differentiation, supporting the hypothesis that STAU2 loss promotes a shift from proliferation toward early differentiation. **C,D.** GSEA curves show upregulation of glycolysis **(C)** and oxidative phosphorildation **(D)** in neuroepithelial cells. **E.** MA plot of differentially expressed genes in neuroepithelial cells. Orange and blue dots represent significantly upregulated and downregulated genes in STAU2 KO, respectively. Green-highlighted genes are miRNA host genes. **E,F.** Cumulative distribution function plots showing the change in expression of predicted targets of miR-9-5p **(E)** and miR-218-3p **(F)** in STAU2 KO neuroepithelial cells versus WT neuroepithelial cells. In both cases, miRNA target gene sets show significant downregulation compared to genes not targeted by miRNAs (Kolmogorov-Smirnov test adjusted p-value = 3.04e-05 & 2.97e-04 respectively). **F.** Bar plot showing the difference in the proportion of cells with an active GRN between STAU2 KO and control samples for selected GRNs. Blue and orange bars mark the GRNs more active in control and STAU2 respectively.

### STAU2 KO drives metabolic reprogramming and accelerates neuronal maturation

To further investigate the functional impact of STAU2 KO, we performed Gene Set Enrichment Analysis (GSEA) across individual cell clusters^35^, which revealed both cell-type specific and shared alterations across clusters **(Table S6)**. Across multiple clusters, we observed an overall dysregulation of cholesterol metabolism, RNA metabolism, translation and synaptic signaling (Fig. S7). In several NPC subtypes and astrocyte progenitors an enrichment for gene sets related to cholesterol and steroid biosynthesis pathways, including *Metabolism of steroids*, *Regulation of cholesterol biosynthesis by SREBP (SREBF)*, *activation of gene expression by SREBP (SREBF)*, and *WP_cholesterol biosynthesis pathway in hepatocytes* **(Table S6)**. Notably, cNPCs3 and TAC3+ Interneurons also showed enrichment of the *cholesterol metabolism with Bloch and Kandutsch–Russell pathways*, a hallmark pathway of neuronal sterol synthesis. Importantly, intact cholesterol synthesis is essential during development, as neurons at early stages depend on endogenous cholesterol production for axonal outgrowth, dendritogenesis, and survival^36^.

In neuroepithelial cells, STAU2 KO led to a significant enrichment of oxidative phosphorylation (OXPHOS), electron transport chain activity, ROBO-SLIT signaling, and glycolysis-related pathways including MYC and mTORC1 signaling among others **(Fig. 3B- D, Fig. S8, Table S6)**. Previous studies have shown that a metabolic shift from glycolysis towards OXPHOS is associated to the transition from proliferative neural progenitors to committed neurogenic cells^37,38^, further suggesting that STAU2 KO accelerates neuroepithelial differentiation by enabling an early metabolic shift. Beyond metabolism, GSEA highlighted additional processes related to cell cycle progression, transcriptional regulation, post-transcriptional control, and protein translation, pointing to a broader role for STAU2 in shaping gene expression programs. Of particular note, we observed enrichment of the SLIT2/ROBO signaling pathway, a key regulator of axon migration, across multiple neuronal clusters. In parallel, neuronal clusters such as interneurons, excitatory neurons, and TAC3LJ interneurons, showed enrichment of *neurotransmitter receptors and postsynaptic signal transmission*, *neuronal system*, and *L1CAM interactions* **(Table S6)**. These results are consistent with the hypothesis that STAU2 regulates the timing of neuronal maturation in neuroepithelial cells.

### STAU2 KO alters miRNA activity in neuroepithelial cells

One of the pathways significantly upregulated in the GSEA analysis in neuroepithelial cells is *Alzheimer’s Disease and miRNA effects* **(Table S6).** This pathway contains several genes related to Alzheimer’s Disease (AD) including genes involved in APP processing, TAU, mitochondrial function, proteasome and autophagy, among others, which are in part regulated by miRNAs differentially expressed in AD **(Fig. S9)**. In the differential expression analysis of neuroepithelial cells, we found that several of the top DEGs corresponded to miRNA host genes –genes from which mature miRNAs are transcribed and processed **(Table S7, Fig. 3E** green dots). These include, among others, C1orf61/MIR9-1HG and SLIT2, host genes of miR-9 and miR-218 respectively, which are well-known regulators of neurogenesis^39^. Although single-cell RNA-seq does not capture mature miRNA levels, we reasoned that changes in host gene expression could affect the relative abundance of mature miRNAs, contributing to gene expression changes in neuroepithelial cells. To test this hypothesis, we retrieved predicted miRNA targets from TargetScan 8.0 database^40^ and compared the expression of miRNA target genes versus non-target genes. Remarkably, target genes of 35 out of 48 miRNAs associated with differentially expressed host genes were significantly downregulated in STAU2 KO neuroepithelial cells (FDR < 0.001**, Table S7**). These included targets of miR-9-5p and miR-218-3p **(Fig. 3F, G),** as well as the miR- 106b∼25 cluster embedded in the MCM7 genomic locus, which is known to regulate adult neural stem and progenitor cell proliferation and neuronal differentiation through TGFB/IGF- insulin pathways^41^.

Importantly, downregulation of miRNA target genes occurred regardless of whether the host transcript was upregulated and downregulated, probably reflecting that STAU2 impact on miRNA processing depends on the location of its binding sites relative to the pre-miRNA hairpin.

We additionally performed qPCRs to validate the changes in the expression of host genes and their effect on the expression of mature miRNAs. We confirmed changes in the expression of *SLIT2* and *MIR9-1HG* in D11 cultures as well as the expression of mature miR-9 **(Fig. S10)**. While both the scRNA-seq and the qRT-PCR show significant changes in the expression of the host genes and miR-9, the direction of these changes is different in these two experiments likely reflecting contributions from multiple early cell population (neuroepithelial cells, early NPCs and early cortical neurons) that cannot be disentangled in bulk. Together, these results suggest that during early neurogenesis, STAU2 is associated with the regulation of miRNAs and repression of their target mRNAs, potentially contributing to altered gene expression in neuroepithelial cells.

### Gene regulatory network analysis identifies ARID3A and CHD2 as downstream mediators of STAU2 function

To better understand the mechanisms driving the accelerated differentiation observed in STAU2 KO cells, we performed Gene Regulatory Network (GRN) analysis using our single-cell transcriptomic data. A GRN is defined as a set of transcription factors and their target genes that interact to regulate specific functions within a cell. We used SCENIC^42^ to infer active GRNs across all cells in our dataset and identified 374 GRNs, many of which show clear cell-type specific activity. For example, NANOG was active in iPSCs, MSX1 in astrocytes, E2F4 in progenitor cells and NEUROD2 in neurons –consistent with previously described roles for these factors^43–46^ **(Fig. S11, Table S8)**.

To assess whether STAU2 KO alters GRN activity in a cell-type manner, we used the binarized output matrix from SCENIC, which indicates whether a specific GRN is considered active or inactive in each cell, to compare the conditions between STAU2 KO and WT **(Table S9)**. STAU2 KO neuroepithelial cells exhibited significantly decreased activity of regulons driven by chromodomain Helicase DNA Binding Protein 2 (CHD2) and AT-Rich Interaction Domain 3A (ARID3A), both of which are transcription factors (TFs) involved in stem/progenitor regulation in different tissues^47,48^. Additionally, we also noticed significant downregulation of several lineage defining TF such as OCT1, PAX6 and GATA4^49–51^ **(Fig. 3H)**. Interestingly, both CHD2 and ARID3A have been previously reported as direct targets of STAU2^21^. CHD2 is a chromatin remodeler that is known to be crucial during development^52^. Dysfunctional CHD2 has been associated with neurodevelopmental disorders such as epileptic encephalopathies^53^. ARID3A is a TF that regulates cell cycle progression and differentiation in various lineages, including hematopoietic and intestinal stem cells^48^. Decreased activity of both TFs increases differentiation in their respective systems^47,48^. In our dataset, we observed that both ARID3A and CHD2 expression and activity were significantly reduced in STAU2 KO **(Table S5, Fig. 3H)**, supporting the idea that STAU2 modulates early neural development by controlling the availability of key transcriptional regulators.

Given that we previously found that STAU2 regulated miRNA host genes and potentially miRNA function, we tested if those miRNAs were additionally contributing to the downregulation of specific GRNs in neuroepithelial cells. Our analyses showed that there was a significant overlap between genes included in downregulated GRNs controlled by ARID3A and CHD2 in STAU2 KO and those targeted by miR-9-5p (Chi-square test p-value = 3.365e-06, **Table S10**). miR-9 host gene is significantly differentially expressed in STAU2 KO neuroepithelial cells, indicating a coordinated regulation of gene expression by STAU2 both at the transcriptional and the post-transcriptional level. Together, these findings position CHD2 and ARID3A as likely mediators of the accelerated neurogenesis phenotype observed in STAU2 KO cells. When looking at genes in upregulated GRNs, we found no significant changes compared to cell-type specific GRNs (**Table S10**, adjusted p-value < 0.001). These results strongly indicate STAU2 regulates gene function both transcriptionally via GRNs and post-transcriptionally through miRNA activity.

### STAU2 loss-of-function significantly affects early neurogenesis and cortical organoid development

The previous computational analyses predicted that STAU2 drives metabolic changes and accelerates neuronal differentiation of neuroepithelial cells. To experimentally validate these predictions, we performed immunofluorescence staining to quantify the proportion of mature neurons at D11 and D25, time points corresponding to neuroepithelial and early NPC stages respectively **(Fig. S5A & S6)**. At D11, STAU2 KO cultures exhibited a significant increase in the number of MAP2+ and TBB3+ cells compared to wild-type controls, indicating that neuronal differentiation is initiated earlier in the absence of STAU2 **(Fig. 4 A, B & Table S11)**. Similarly, by D25 the number of MAP2+ cells remains higher in STAU2 KO cells **(Fig. 4C, D)**, consistent with an accelerated neuronal differentiation phenotype.

**Figure 4.**
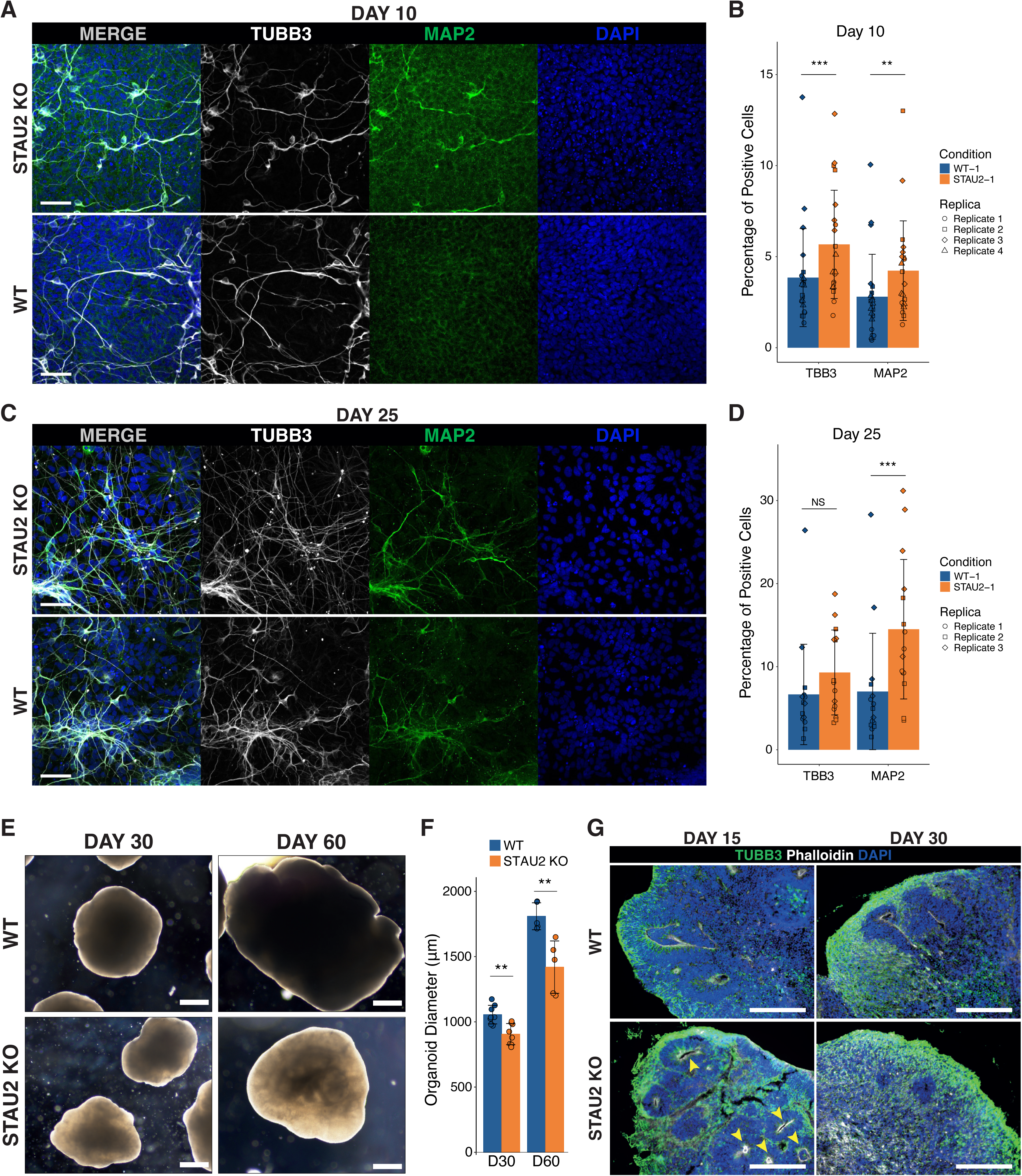
STAU2 KO leads accelerated neural differentiation at the expense of the progenitor pool. **A.** Immunostaining of βIII-Tubulin (TUBB3) and MAP2 cells at D11 of differentiation in STAU2 KO and WT cells. **B.** Quantification of TUBB3+ and MAP2+ cells at D11 in STAU2-KO and WT cells. Colored bars show the average number per condition +/- SD. Measurements obtained from independent biological replicates are highlighted with specific point marks. Comparison between STAU2-KO and WT shows significantly increased neuronal populations in STAU2 KO, suggesting precocious neurodifferentiation (** p-value < 0.01; *** p-value < 0.001 Non-parametric Two-Way ANOVA test). **C.** Immunostaining of TUBB3 and MAP2 cells at D25 in STAU2 KO and WT cells. **D.** Quantification of TUBB3+ and MAP2+ cells at D25 shows a significant increase in the number of MAP2+ cells (*** p- value < 0.001). **E.** Brightfield images of STAU2-1 KO and WT-1 cortical organoids at D30 and D60. STAU2 KO organoids are smaller in size (scale bar: 100 µm). **F.** Quantification of organoid diameter at D30 and D60 reveals a significant reduction in STAU2 KO organoid size compared to WT (** p-value < 0.01; Non-parametric Two-Way ANOVA test). **G.** Immunostaining of cortical organoids stained for TBB3 (green, neuronal marker), DAPI (blue, nuclei), and Phalloidin (white, labeling actin-rich neural rosettes) at D15 and D30 (scale bar: 100 µm). STAU2 KO organoids show increased neuronal differentiation and larger, more prominent neural rosettes compared to WT.

The increase in the number of early neurons, together with the results of the computational analyses, which identified an upregulation in pathways associated with neurogenic metabolic activation^54–56^, led us to hypothesize that STAU2 loss promotes premature neuronal differentiation at the expense of the progenitor pool, reducing the stem cell population in the cultures. To investigate this hypothesis, we generated cortical organoids from the two independent STAU2 KO cell lines and their isogenic controls. Across both cell lines, STAU2 KO organoids were consistently smaller compared to the WT counterparts at early and late stages of development **(Fig. 4E and Fig. S12A)**. Quantification of organoid diameter at D30 revealed a reduction in size in STAU2 KO samples. This difference became even more significant by D60, suggesting impairments in tissue growth, likely resulting from impaired progenitor expansion or early progenitor depletion **(Fig. 4E, F & Fig. S12A, B)**. Immunofluorescence analysis at D15 showed that both WT and STAU2 KO organoids formed rosette-like structures characteristic of early neuroepithelial organization. However, neuroepithelial loops were smaller in KO organoids **(Fig. 4G & Fig. S12C, yellow arrows & Fig. S13)**. Importantly, STAU2 KO organoids displayed an unexpectedly higher number of MAP2^+^ neurons (MAP2 staining, green) at early developmental stage indicating premature neuronal differentiation, despite reduced size. By D30, MAP2 signal intensities became more comparable between WT and STAU2 KO organoids, consistent with progenitor pool exhaustion in KO condition **(Fig. 4G & Fig. S12C)**. Together, these findings demonstrate that STAU2 is essential for maintaining the balance between progenitor self-renewal and neuronal differentiation in human cortical development. Loss of STAU2 impairs progenitor expansion and early tissue organization, leading to premature neuronal differentiation and reduced growth in cortical organoids.

## Discussion

RBPs are pivotal regulators of gene expression, orchestrating various co- and post- transcriptional processes that influence cell fate decisions during development. Among these, the double-stranded RBP STAU2 has emerged as a key player in neural development. In this study, we investigated the expression and function of STAU2 during human brain development by knocking out the expression of STAU2 in iPSCs and evaluating its impact on neural and glial differentiation using single-cell transcriptomics. Single-cell transcriptomic profiling across multiple time points revealed that loss of STAU2 significantly impacts neuroepithelial cells, the earliest neural progenitors at different levels, and other NPC clusters to a lower extent. These findings extend previous work on mouse embryonic cortex^16,17^ by demonstrating that STAU2 plays a critical role in regulating the maintenance and differentiation of neural progenitor cells during early human corticogenesis. As gene expression changes were most pronounced at this early stage, we focused on the role of STAU2 in neuroepithelial cells. Mechanistically, our data reveal that STAU2 controls early lineage decisions through multiple, interconnected regulatory layers. First, metabolic rewiring emerges as an early hallmark of STAU2 loss. STAU2 KO neuroepithelial cells showed a striking upregulation of both glycolytic and OXPHOS pathways. Neural stem and progenitor cells generally rely on glycolysis to support proliferation under hypoxic conditions, whereas neuronal differentiation requires increased mitochondrial respiration and OXPHOS^54–56^, providing the energy required for the function and maintenance of mature neurons. Not only were these pathways enriched in STAU2 KO neuroepithelial cells, but we also identified additional molecular pathways involved in the self-renewal and differentiation of neural stem cells into mature neurons, including the mechanistic Target of Rapamycin (mTOR), and hypoxia signaling^57,58^ **(Fig. 3B)**. The enrichment of these pathways suggests a metabolic shift typically associated with early neuronal differentiation, which indicates that these cells are undergoing a premature transition toward a neuronal metabolic profile. This interpretation aligns with the observed acceleration of neuronal differentiation in STAU2 KO cultures, where we detected an elevated proportion of MAP2⁺ neurons compared to control (WT) conditions **(Fig. 4A-D)**. This molecular shift also aligns with phenotypic changes observed in STAU2 KO 3D organoid systems, which display premature expression of neuronal markers, loss of laminar organization, and reduced size, features that suggest early progenitor pool exhaustion, when symmetric divisions should increase the number of cells of the stem cell niche. Previous works show that Stau2 depletion speeds differentiation of RGCs with an early production of IPCs and neurons, perturbing cortical development in mouse^16,17^. In this regard, we also observed a consistent enrichment of cholesterol and steroid biosynthesis pathways across several progenitor subtypes (cNPCs and astrocyte progenitors) and interneuron clusters. This is highly relevant, as the brain contains ∼25% of the body’s cholesterol, which cannot cross the blood–brain barrier and therefore must be synthesized locally^36^. Developing neurons rely on *de novo* cholesterol synthesis for axonal outgrowth, dendritogenesis, and survival, while glial cells later contribute to sterol homeostasis. These findings highlight cholesterol metabolism as a key downstream process perturbed by STAU2 loss. We propose that STAU2 acts to maintain the progenitor state by regulating gene expression programs associated with metabolic shifts and neuronal maturation. Its loss destabilizes this balance, leading to early activation of differentiation-associated pathways and metabolic remodeling characteristic of neurogenesis. In fact, prior studies have identified specific clusters of RBPs that respond to metabolic changes, particularly oxygen availability. Proteins such as HuR, PCBP1 and hnRNP A2/B1 enhance the translation of hypoxia-responsive proteins involved in glycolysis, such as enolase 1 (ENO1), aldolase A and lactate deshydrogenase A (LDHA)^59^.

Second, among the differentially expressed genes we identified several miRNA host genes whose mature miRNAs are involved in neuronal differentiation. The list of miRNA host genes **(Table S7)** includes several previously validated STAU2 targets, such as MIR9-1HG and SLIT2^7,17,21^. Gene expression changes in these miRNA host genes also impacted their corresponding mature miRNAs, as seen for miR-9, and were associated with a concordant downregulation of their validated mRNA targets in STAU2 KO cells **(Fig. 3F, G, Table S11)**. Both miRNAs are well-established regulators of neurogenesis: miR-9 is a highly conserved, neuronally enriched miRNA essential for coordinating neurogenesis, promoting neuronal differentiation, and restricting progenitor proliferation by targeting key regulators such as TLX, SIRT1, and BACE1^39,60–63^. miR-218, hosted within the SLIT2 locus, also regulates neuronal radial migration and morphogenesis during cortical development, in part by repressing Satb2 in mice^64^. Additionally, we detected changes in the MCM7 locus encoding the miR-106b∼25 cluster, also affected in STAU2 KO cells, which has been shown to regulate neural stem cell and progenitor proliferation as well as TGFβ/IGF signaling during differentiation^41^. These data support a model in which STAU2 buffers miRNA-mediated transcript repression in neuroepithelial cells, including miR-9 activity, thereby preserving a progenitor state and preventing premature differentiation. This model is consistent with previous evidence showing that interactions between RBPs and miRNAs fine-tune gene expression in neural development as observed for Musashi1 (Msi1) and Elavl1^39,65–67^.

Third, in addition to transcriptomic and morphological changes, the GRN analysis suggests that STAU2 also modulates neurogenesis through selective post-transcriptional control of key developmental regulators. We identified reduced activity of transcription factors CHD2 and ARID3A, which were previously validated as targets of STAU2^21^, as potential drivers of the accelerated differentiation phenotype. Importantly, the downregulation of CHD2 and ARID3A in STAU2 KO neuroepithelial cells may function upstream of the metabolic shift toward oxidative phosphorylation and glycolysis. CHD2, a chromatin remodeler essential for maintaining radial glial self-renewal, has been shown in mouse cortex to preserve progenitor identity by sustaining chromatin accessibility and balanced transcriptional programs. CHD2 mediates these functions binding to the genomic region of repressor 1-silencing transcription factor (REST) and its loss leads to premature neuronal differentiation^47,68^. Although direct studies in neural contexts are limited, CHD2 is known more broadly to regulate transcriptional programs governing cell cycle and metabolic genes, suggesting that its suppression could release metabolic genes from epigenetic constraint^47,68^. Similarly, ARID3A is a DNA-binding protein implicated in lineage timing and chromatin remodeling. In stem cell systems, ARID3A loss accelerates differentiation by disrupting transcriptional networks, and it interacts with histone deacetylases, linking it to epigenetic control of gene expression^69^.

We propose that in WT cells, STAU2 stabilizes CHD2 and ARID3A mRNAs, preserving a progenitor-specific epigenetic landscape that restrains metabolic gene activation. Loss of STAU2 downregulates these factors, loosening chromatin control and enabling early activation of metabolic (MYC, glycolysis, OXPHOS) programs, which, in turn, prime premature neurogenesis and maturation (Fig. 5).

**Figure 5.**
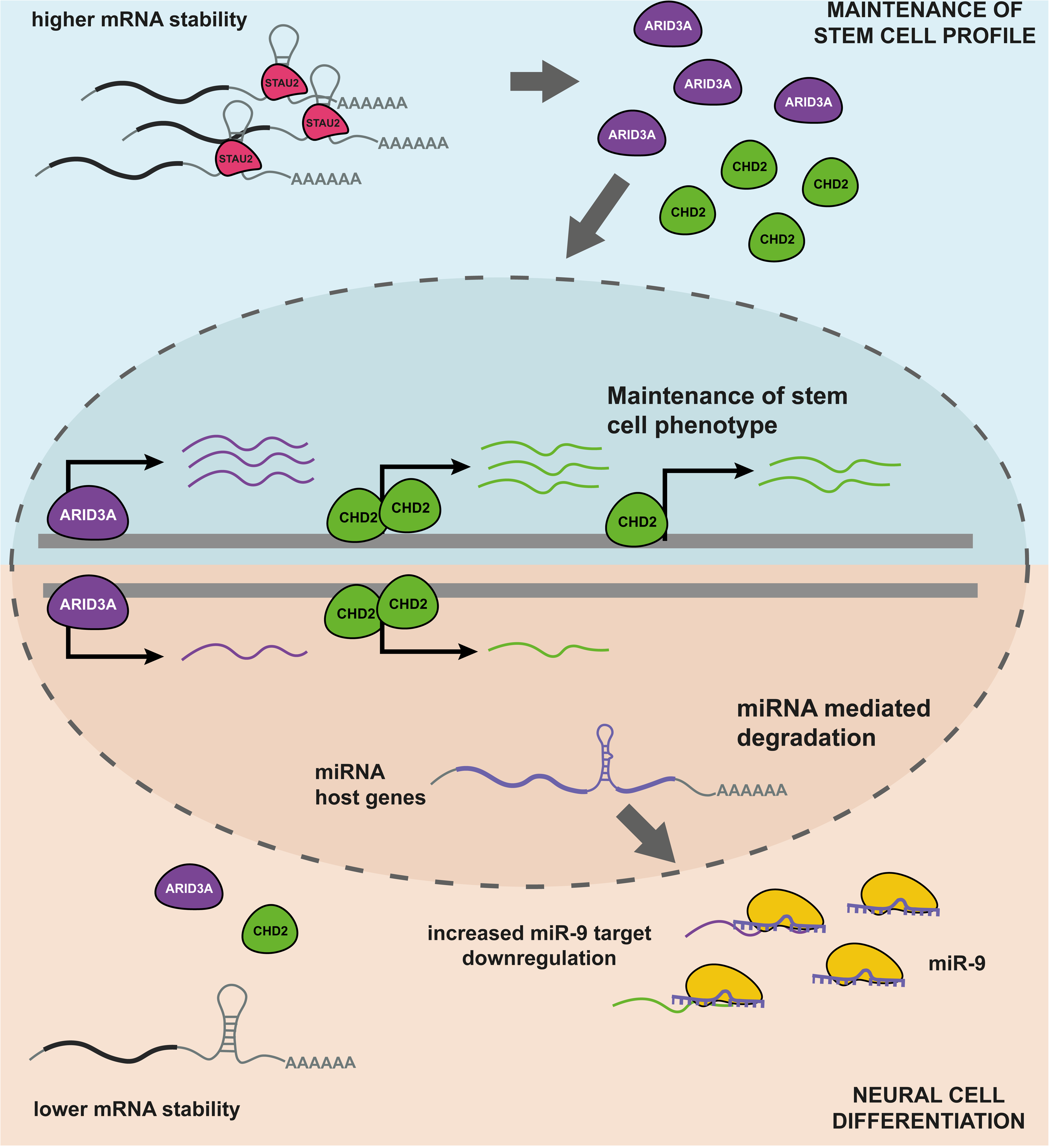
Proposed model for STAU2 regulation of neurogenesis at the neuroepithelial level. In WT conditions (blue background), STAU2 stabilizes the mRNAs of ARID3A and CHD2, which activate transcriptional programs maintaining the stem cell phenotype. Upon KO, downregulation of ARID3A and CHD2 mRNAs leads to a reduced activity of their GRNs and the loss of stem cell phenotype. The higher activity of miR-9, which targets the genes regulated by ARID3A and CHD2 provides additional control to downregulate ARID3A & CHD2 GRNs and promote the neural differentiation.

Interestingly, we observed a significant overlap between miR-9-5p and miR-218 targets and downregulated GRNs in neuroepithelial cells (**Table S10**, Chi-square test p-value = 1.64e-08 and p-value = 2.55e-03 respectively). This suggests that STAU2 controls miRNA function and transcription factor activity in a coordinated manner to regulate the genetic programs during early neurogenesis. Together, our findings position STAU2 as a central node in post-transcriptional regulation during human corticogenesis, operating at the intersection of miRNA activity, transcription factor output, epigenetic state, and metabolic control. This model supports a dual role for STAU2, one as gatekeeper of metabolic activation and a post-transcriptional regulator of epigenetic programs, coordinating multiple regulatory layers to ensure the timely progression and proper identity of emerging neural cell types. Future studies should explore whether STAU2 plays similar roles in other human stem cell populations and how its dysregulation may contribute to neurodevelopmental disorders.

## Online Methods

### Generation of functional STAU2 KO cell lines

STAU2 KO iPSC cell lines (STAU2-1 & STAU2-2) were generated using CRISPR/Cas9- mediated genome editing targeting a conserved double-stranded RNA-binding domain (dsRBD2) shared across major STAU2 isoforms. The exons 7 and 8 of STAU2 were deleted by using two single guide RNAs (sgRNAs) flanking this region **(Fig. 1A)**. sgRNAs were designed for high on-target efficiency and minimal off-target potential (IDT, Alt-R® CRISPR- Cas9 sgRNA Design Tool). Shortly, iPSCs were nucleofected with the RNP Complex: Alt-R® CRISPR Cas9 sgRNA + Cas9 Protein (IDT) using the Amaxa 4D-Nucleofector™ system (Lonza) and the electroporation protocol CA-137. Nucleofected cells were then plated onto 48-well plates containing pre-warmed mTeSR1 medium (StemCell Technologies) supplemented with 10 µM of Y-27632 ROCK inhibitor (StemCell Technologies, #72304) to promote survival. Once cultures reached 80% confluency, cells were dissociated into single cells. 1,000 cells were plated on top of irradiated human fibroblast feeders with mTeSR1 complete medium on 15 cm dishes and maintained for 10-14 days until colonies were large enough to be screened. Colonies were manually picked and genotyped by PCR and Sanger sequencing **(Fig. 1A)**. Colonies with the desired genotype were isolated, expanded, and cryopreserved. STAU2 deletion was verified by Western Blot and qPCR **(Fig. 1B, C)**. The STAU2 KO generated cell lines were karyotyped to check genomic integrity **(Fig. S1)** and tested for pluripotency capability **(Fig. 1D)**.

### iPSC cell culture and differentiation

STAU2 KO iPSCs (STAU2-1 and STAU2-2) and isogenic wild-type control iPSC lines (WT1 and WT2) were cultured and differentiated toward neural lineages following the protocol described previously^29^. For single-cell sequencing, cells differentiated from STAU2-1 cell line were harvested at five developmental stages corresponding to days 0, 11, 25, 55, and 70 of differentiation and fixed in methanol using our previously described protocol^70^ for transcriptomic profiling. Differentiations were carried out in two independent batches, each including one biological replicate per genotype (STAU2 KO and wild-type control), yielding a total of four differentiations. At each timepoint, cells were collected for single-cell transcriptomic analysis using Drop-seq, as described previously^71^.

### Cell dissociation and sample preparation

Cells collected at the five differentiation stages (D0, D11, D25, D55, and D70) were dissociated into single-cell suspensions following protocols optimized for single-cell RNA sequencing (scRNA-seq), as previously described^70^. Briefly, early-stage cultures (D0 and D11) were incubated with accutase (StemCell Technologies) for 5–8 minutes at 37°C. For later timepoints (D25, D55, D70), cells were dissociated using a papain-based enzymatic mix (1:1 ratio of papain and accutase) prepared using the PDS Kit (Worthington Biochemical Corporation, #LK003176), with incubation times ranging from 25 to 35 minutes depending on tissue density. Dissociated cells were resuspended in neural maintenance medium supplemented with 10 µM of Y-27632 and 0.033 mg/ml of DNase I (Worthington Biochemical Corporation, #LK003170). Single-cell suspensions were passed through a 40 µm cell strainer (Pluriselect Life Science, #43-10040-60) to remove cell aggregates, centrifuged at 150 x g for 3 minutes at RT and washed 3 times with 0.4 mg/ml BSA in DPBS. After cell counting, 2.5 x 10^6^ cells were fixed in methanol following the most recent recommendations to perform single-cell transcriptomics on iPSC-derived neural cells^70^. Only samples with cell viability higher than 85% before preservation were included in the study. Prior to encapsulation, methanol-fixed cells were thawed on ice, pelleted, and re-hydrated in 1 ml of DPBS containing 0.01% BSA, 0.2 U/µl of RNase inhibitor (Takara Bio, #2313A), and 1 mM DTT (Merck Sigma-Aldrich, #D0632). Before single-cell encapsulation, cells were filtered again through a 40 µm strainer to minimize doublet formation.

### Single-cell transcriptomics

Drop-seq methodology was performed using the NADIA microfluidic instrument (Dolomite Bio, #3200590) following the manufacturer’s protocol^72^ without modifications. Briefly, 250 µl of a single-cell suspension (300,000 cells/ml) was co-flowed with 250 µl of Macosko oligodT beads (ChemGenes Corporation, #Macosko-2011-10(V+)) at 600 beads/µl, previously washed and resuspended in lysis buffer (6% w/v Ficoll PM-400, 0.2% v/v Sarkosyl, 20 mM EDTA, 200 mM Tris pH 7.5 and 50mM DTT in nuclease-free water). The RNAs captured by the oligodT are immediately reverse transcribed using Maxima H Minus Reverse Transcriptase (Thermo, #EP0751) after emulsion breakage. Then, beads are incubated with exonuclease I (New England Biolabs, #174M0293L). Beads were counted and resuspended at 400 beads/µL in nuclease-free water. PCR amplification was carried out in aliquots of 4,000 beads per reaction and amplified for 11 PCR cycles. Amplified cDNA was purified using AMPure XP Beads (Agencourt, #A63881) at 0.6:1 ratio to sample. cDNA concentration was quantified with Qubit dsDNA HS Assay (Thermo, #Q32851), and fragment size was measured by a 4200 TapeStation System (Agilent, #G2991BA). Library construction was performed using the Nextera XT DNA Library Prep Kit (Illumina, #FC-131-1096) starting from 600 pg of amplified cDNA to tagment, tag and amplify. The resulting library was purified with 0.6:1 AMPure XP Beads to sample and fragment size was determined using a 4200 TapeStation System and quantified with Qubit dsDNA HS Assay. Pooled libraries (final concentration:1.8-2.1 pM) were sequenced on Illumina NextSeq 550 sequencer using Nextseq 550 High Output v2 kit (75 cycles) (Illumina, #20024906) in paired-end mode. Read 1 (20 bp) was sequenced using the custom primer Read1CustSeqB^32^ (cell barcode and UMI), Read 2 (64 bp) captured the transcript sequence, and an 8 bp i7 index was used for sample demultiplexing.

### Immunocytochemistry

Cells were fixed at selected time points during neural differentiation using a stepwise paraformaldehyde (PFA) fixation protocol: sequential incubations in 2%, 3%, and 4% PFA (each for 5 minutes at room temperature (RT) and washed three times for 5 min with PBS. Permeabilization and blocking were performed simultaneously with 0.5% Triton X-100 and 6% Normal Donkey Serum (NDS; Merck Sigma-Aldrich #D9663) in Tris-buffered saline (TBS) for 1 hour. Primary antibody incubation was performed in 0.1% Triton X-100 and 6% NDS in TBS (TBS++) for 48 hours at 4°C (see **Table S12** for antibody list and concentrations). After incubation, samples were washed three times, 5 min with TBS. Secondary antibodies, also diluted in TBS++, were incubated overnight at 4°C **(Table S12)**. Following incubation, samples were washed three times with TBS. Nuclear staining was performed by a 10 min incubation at RT with DAPI (4,6-diamidino-2-phenylindole) at 1:10,000 dilution in TBS. Slides were mounted with PVA: DABCO and stored at 4°C until imaging.

Fluorescence imaging was performed using a Carl Zeiss LSM 880 spectral confocal laser scanning microscope (Carl Zeiss Microscopy GmbH, Jena, Germany) equipped with a multiline argon laser (458, 488, and 514 nm), 405 and 561 nm diode lasers and 633 nm He/Ne laser (Centres Científics i Tecnològics, Universitat de Barcelona, Bellvitge Campus, Barcelona, Spain). Images were taken using a 25x oil immersion objective (0.8 NA), single image resolution of 1024 X 1024 pixels, pinhole aperture adjusted to 1Airy unit 3x3 tiles imaging.

Image analysis was performed using Fiji/Image J (Wayne Rasband, NIH, USA). For nuclear positive staining quantification, we run a customized macro script. Nuclear segmentation was performed using the StarDist2 plugin. Detected ROIs were filtered by area to exclude debris and cell clusters (thresholds: <25 and >700 arbitrary units). For immunopositive signal quantification, the channel of interest was binarized, and each ROI was applied to extract overlapping signal. A nucleus was considered positive when >10% of its area overlapped with the binary mask. Output files included ROI coordinates, total DAPI counts, positive cell counts, and percent of counts relative to total DAPI.

### Quantitative real-time PCR (qRT-PCR)

Total RNA was extracted using the Maxwell® RSC simplyRNA Cells Kit (Promega) following the manufacturer’s instructions. 1 µg of total RNA was reverse transcribed into cDNA using the Transcriptor first-strand cDNA synthesis kit (Roche, cat. no. 04896866001), following the manufacturer’s instructions. qRT-PCR was performed using 50 ng of cDNA per reaction, run in duplicates or triplicates on a **L**ightCycler® 480 System (Roche) with 480 SYBR Green I Master Mix, (Roche, #04707516001). Non-template controls and non-reverse transcription controls were included for each probe to rule out contamination and ensure specificity.

Reactions were run under the following thermal cycling conditions: 15s at 95°C for denaturation and 1 min at 60°C for annealing and extension, for 40 cycles. CT values were measured, and relative gene expression levels were calculated using the 2^–ΔΔCT method. Human PPIA was used as a housekeeping gene for normalization unless stated otherwise in the figure legend. Primer sequences are provided in **Table S13**.

### TaqMan Assay

Mature miR-9 was measured using a standard TaqMan Assay (Applied Biosystems, #4440887, assay ID #000583) performed according to the manufacturer’s instructions. RPL21 (TaqMan assay ID #001209) and RNU58B (TaqMan assay ID #001206) were used for normalization. qPCR reactions were performed using TaqMan™ Universal Master Mix II, no UNG (ThermoFisher Scientific, # 440047) in technical triplicates. Relative miRNA expression calculated using the 2^–ΔΔCT method.

### scRNA-seq Data pre-processing

Drop-seq tools v2.4.0 pipeline^73^ was used to generate the transcript count matrices from the processed scRNA-seq libraries. Briefly, we first generated the index and annotation files for the GRCh38.93 assembly of human genome with ENSEMBL version 93^74^. We merged the FASTQ files with paired-end reads into unaligned BAM files using Picard tool FastqToSam v2.18.25^75^. Adapter sequences and polyA tails were trimmed at the 5’ and 3’ ends respectively, using the Drop-seq toolkit with default parameters. Trimmed reads were aligned to hg38 Human Genome using STAR version 2.7.8a^76^. The annotation files generated previously were used to identify the reads overlapping genes. To correct cell barcode errors, we used the DetectBeadSubstitutionErrors and DetectBeadSynthesisErrors programs with default parameters. We estimated the number of cells captured during single-cell encapsulation using a knee plot. The number of uniquely mapped reads assigned to the top N barcodes was used as input, with N being five times the expected number of cells. Finally, we used the function DigitalExpression to generate the transcript count matrices with the cell number estimate and the parameters LOCUS_FUNCTION_LIST=INTERGENIC LOCUS_FUNCTION_LIST=INTRONIC LOCUS_FUNCTION_LIST=UTR LOCUS_FUNCTION_LIST=CODING.

### scRNA-seq quality control

We used Seurat version 4.1.1^77^ to analyze the scRNA-seq datasets. First, Seurat objects were generated for each sample. Filtering out low-quality cells was done on a per-sample basis as recommended previously^78^. First, cells with gene number and UMI count < 200 were discarded. In D0, D11 and D25 samples, cells were retained if the mitochondrial percentage (percent.mt) was lower than 5%. Cells from D55 and D70 samples were retained if percent.mt < 8%. In the subsequent filtering steps, the cells are first tagged for discard and cells failing any of the filtering steps are discarded at the end. Such an approach prevents the order of filtration steps from affecting which cells are removed. Cells in the top 1% of UMI count in each sample were marked. Subsequently, doublets were identified using DoubletFinder^79^ and marked for discard. Lastly, a linear model was fitted between log- transformed gene number and UMI count of cells in the combined Seurat object, and cells with residuals less than -0.7 or greater than 0.3 were marked. Cells marked in at least one step are discarded. Subsequently, genes expressed in less than three cells were also discarded. The final Seurat object consists of 29368 cells and 27138 genes.

### Clustering and cell type identification

We used the SCTransform function to normalize data and to regress out the effect of confounders (gene number, UMI count, mitochondrial and ribosomal percentage, cell cycle scores). For each experiment, we identified 5000 Highly Variable Genes (HVGs). HVGs found in at least three of the differentiations or found in two differentiations of the same condition (KO or WT) were used to get 3000 integration features to perform Canonical Correlation Analysis. After integrating the datasets, we ran Principal Component Analysis on the integrated Seurat object to identify 100 Principal Components (PCs) and visually inspect the Elbowplot generated to study the amount of variability explained by each PC. The first 18 PCs were selected to generate the kNN graph and UMAP plot. The package clustree^80^ was used to understand how cells are assigned to clusters with resolution 0.1 to 1.0. After manual inspection, we chose the resolution of 0.6 for downstream analysis. We used the function FindAllMarkers of Seurat v 5.0.1^31^ to identify cell-type specific marker genes with the following parameters: only positive markers with min.pct = 0.25 and logfc >= 0.25. Significant markers (Wilcoxon rank-sum test; FDR < 0.05) were used along with markers from literature to annotate the clusters.

### Cell Composition Analysis

We used the scCODA package^32^ to study changes in cell composition between KO and WT cells at different timepoints. First, we counted the number of cells in KO and WT for each cluster in each sample (each time point) **(Table S4)**. Based on dispersion and cell number, we manually selected a reference cluster that had a low variance between conditions for each of the time points. We employed a linear model with Condition and Batch (∼ Condition + Batch) to find changes in cell composition considering both factors. scCODA uses Hamiltonian Monte Carlo (HMC) simulations to detect changes in cell composition of clusters for a condition. If the output score for a cluster is greater than 0, it is enriched and if it is less than 0, it is depleted. Due to stochasticity of cell composition analysis with scCODA, we run the HMC 10 times and if enrichment or depletion is concordantly detected in at least 6 runs, we considered it a significant change in cell composition for that cluster.

### Differential Expression Analysis

We used the FindMarkers function with the MAST test from Seurat version 5.0.1^31^ to identify Differentially Expressed Genes (DEGs) between KO and WT samples within each cluster. Only genes expressed in at least 10% of cells in either condition (KO or WT) and |log2FoldChange| > 0.1 were considered. DEGs with |log2FoldChange| > log2 (1.5) and FDR < 0.05 were considered significant. Confounders such as batch, ribosomal and mitochondrial percentage, number of genes and UMIs detected were included as latent variables in MAST analysis.

### Gene Set Enrichment Analysis

Genes were ranked by signed log_10_ p-value so that more significantly upregulated or downregulated genes were at the beginning and the end of the list respectively. This ranked list was used to perform the pre-ranked gene set enrichment analysis (GSEA) using the GSEA-P application^35^ with gene sets from the Molecular Signature Database (MSigDB) ^81–83^. We considered significantly gene sets enriched or depleted with an FDR < 0.05.

To better visualise enriched genesets, we used the rrvgo package^84^. Briefly, based on gene overlap, we grouped genesets from Hallmark, Reactome and WikiPathways collections in (MSigDB) ^81–83^. Pairwise similarities between genesets were quantified using the Jaccard Index, given by the number of shared genes divided by the total number of unique genes across the two sets. We constructed a similarity matrix based on these indices. Genesets with similarity of at least 0.7 were clustered using a custom implementation of the reduceSimMatrix function from the rrvgo package^84^, modified to work without GO-specific dependencies. The results were visualised by multidimensional scaling (MDS) of the Jaccard distance matrix (1-similarity). FDR scores and setSize from GSEA results were incorporated to color and scale point sizes in the plot.

### Gene Regulatory Network inference

The Python implementation of SCENIC (Pyscenic version 0.12.1)^42^ was used to infer the GRNs. The reference files for transcription factors (allTFs_hg38.txt) and databases (hg38_10kbp_up_10kbp_down_full_tx_v10_clust.genes_vs_motifs.rankings and hg38_500bp_up_100bp_down_full_tx_v10_clust.genes_vs_motifs.rankings) were obtained from https://resources.aertslab.org/cistarget. To perform the SCENIC analysis, we first input the raw counts from the filtered Seurat object to the ‘grn’ GRNBoost2 algorithm with default parameters to generate the co-expressing regulatory modules. These modules were pruned using the ‘ctx’ function with cisTarget masking dropouts, only retaining modules enriched for candidate targets of a transcription factor. These modules were grouped into GRNs, each represented by a transcription factor. Finally, the AUcell program was used to calculate the activity of each GRN in each cell of the input count matrix. SCENIC output also gives a binary matrix showing whether a GRN is active (on) or inactive (off) in a specific cell based on adaptive thresholding of AUC values. We use the Fisher’s exact test to identify differentially active GRNs (p.adj < 0.001) by comparing the binary matrices of STAU2 KO and WT cells in each cell type. We also calculate the percentage of ON cells in STAU2 KO and WT for each GRN in a cluster.

### miRNA Analysis

We identified miRNA host genes by comparing DEGs of the neuroepithelial cell cluster with the HGNC microRNA Host Genes database^85^. We found the miRNA species associated with differentially expressed host genes using the ENSEMBL browser (release 112)^86^. We obtained the miRNA 7-mer targets from TargetScan 8.0 database^40^. We first generated a background set composed of genes expressed in neuroepithelial cells that were not targeted by any miRNA linked to differentially expressed host genes. For each miRNA, we used a Kolmogorov-Smirnov test to check for differences in the cumulative distribution of avg_log2FC between target genes and background genes. P-values were adjusted for multiple testing using Benjamini-Hochberg correction. Gene names were converted to ENSEMBL gene ids where necessary for comparison across databases.

### Overlap of miRNA targets and genes of GRNs

We identified genes from ARID3A and CHD2 GRNs and checked how many were 7-mer targets of a specific miRNA (miR-9-5p and miR-218-1-3p). Subsequently, we compared this to genes from the remaining GRNs using a Chi-square test to compare these distributions **(Table S10)**. We performed a similar analysis overlapping miRNA targets with genes from all significantly downregulated GRNs (P.adj < 0.001 and Odds Ratio < 1) in Neuroepithelial cells and compared it to genes from upregulated GRNs (P.adj < 0.001 and Odds. Ratio > 1) or GRNs not enriched in either condition.

### Organoid generation

The organoids were generated using forebrain protocols following a modified version of Walsh et al.^87^. Briefly, after dissociation into a single-cell suspension using TrypLE, 9,000 cells were seeded per well in 96-well plates with 100 µl of E8 medium containing 50 µM ROCK inhibitor. The next day, 80% of the medium was replaced with E6 medium supplemented with 0.1 µM LDN, 5 µM XAV, and 10 µM SB. On day 5, the medium was replaced with E6 medium supplemented only with LDN and SB (dual SMAD inhibition).

Between days 7 and 9, liquid embedding of the organoids was performed using 2% Matrigel (Corning, 356234) in organoid differentiation medium. This medium consisted of a 1:1 mixture of DMEM/F12 and Neurobasal, supplemented with 1× N2 supplement, 1× B27 - vitamin A supplement, insulin, 2-ME solution, GlutaMAX, MEM-NEAA, and LIF. The embedded organoids were cultured in ultra-low-attachment 6-well plates on an orbital shaker (80 rpm) until day 15.

Subsequently, the medium was replaced with organoid maturation medium, consisting of a 1:1 mixture of DMEM/F12 and Neurobasal, supplemented with N2 supplement, B27 + vitamin A supplement, insulin, 2-ME solution, GlutaMAX, MEM-NEAA, sodium bicarbonate, vitamin C solution, chemically defined lipid concentrates, BDNF, GDNF, and cAMP. Live imaging of the organoids was performed using a Nikon Eclipse Ts2 microscope.

### Immunostaining of brain organoids

For immunostainings, organoids were washed three times with PBS and fixed in 4% paraformaldehyde for 20 to 60 min (depending on the organoid size) at 4 °C, then washed with PBS three times for 10 min each. The tissue was incubated in 40% sucrose (in PBS) until it sunk (overnight) and then embedded in 13%/10% gelatin/sucrose. Frozen blocks were stored at -80 °C, prior to cryosections. 10-12 μm sections were prepared using a cryostat. Sections were incubated with warm PBS for 10-15 min in order to remove the embedding medium and then fixed for additional 10 min with 4% PFA, washed three times with PBS and blocked and permeabilized in 0.25% Triton-X, 5% normal goat serum in PBS for 1 h. Sections were first incubated with primary antibodies **(Table S12)** in 0.1% Triton-X, 5% normal goat serum overnight. They were then washed three times for 10 min each with PBST (0.1% Triton X-100) and incubated with secondary antibodies at RT for 2 h, followed by staining with DAPI (final 1 µg/ml) for 10 min and washed three times with PBST. The images were acquired using a Keyence BZ-X710 (Osaka, Japan) microscope.

## Acknowledgements

The authors thank all present and past members from the Plass Lab and Emma Hammarlund for useful comments and critical discussions. The authors thank Nicole Grieger for her help with iPSC colony genotyping. The authors thank IDIBELL’s histology facility and especially Lola Mulero and José A. Llamas for performing STAU2 KO iPSC characterization. We also thank the Scientific and Technological Centres (CCiTUB), Universitat de Barcelona, and especially Benjamin Torrejón Escribano for his support and advice on confocal microscopy. This research was funded by research projects from the State R&D Program Research Challenges from the Spanish Ministry of Science, Innovation and Universities (Project: PID2019-108580RA-I00 funded by MICIU/AEI /10.13039/501100011033 and Project: PID2022-139580OB-I00 funded by MICIU/AEI /10.13039/501100011033 and FEDER and EU). SMFM work was supported by the European Union’s Horizon 2020 research and innovation programme under the Marie Sklodowska-Curie Grant agreement No 101026821. MP work was supported by a Ramón y Cajal contract of the Spanish Ministry of Science, Innovation and Universities (Grant: RYC2018-024564-I funded by MICIU/AEI /10.13039/501100011033 and by “El FSE invierte en tu futuro”). AJG work is supported by the predoctoral program AGAUR-FI ajuts Joan Oró (2024 FI-1 00072), which is backed by the Secretariat of Universities and Research of the Department of Research and Universities of the Generalitat of Catalonia, as well as the European Social Plus Fund. We thank CERCA Program/Generalitat de Catalunya for IDIBELL institutional support.

## Author contributions

SMFM and MP designed the project. AJG and MP designed computational analyses. SMFM, DRM, AG and MP designed the CRISPR KO strategy. AGF and SMFM performed the CRISPR KO in iPSC cells. SMFM, AGF, NCF and RTG performed iPSC cell culture experiments and scRNA-seq experiments. ARW differentiated and characterized organoids. SMFM and MPW performed qRT-PCR experiments. AJG and LM performed computational analyses. SMFM & MP acquired funding and coordinated the work. SMFM, AGF, AG and MP provided supervision. AJG, SMFM and MP wrote the initial draft with input from all authors. All authors reviewed the final manuscript.

## References

1. Taverna, E., Götz, M. & Huttner, W. B. The cell biology of neurogenesis: toward an understanding of the development and evolution of the neocortex. Annu Rev Cell Dev Biol 30, 465–502 (2014).

2. Terreros-Roncal, J. et al. Impact of neurodegenerative diseases on human adult hippocampal neurogenesis. Science (1979) 5163, 1–12 (2021).

3. Moreno-Jiménez, E. P. et al. Adult hippocampal neurogenesis is abundant in neurologically healthy subjects and drops sharply in patients with Alzheimer’s disease. Nat Med 25, 554–560 (2019).

4. Vaid, S. & Huttner, W. B. Transcriptional Regulators and Human-Specific/Primate-Specific Genes in Neocortical Neurogenesis. International Journal of Molecular Sciences 2020, Vol. 21, *Page* 4614 **21**, 4614 (2020).

5. Wilkinson, G., Dennis, D. & Schuurmans, C. Proneural genes in neocortical development. Neuroscience 253, 256–273 (2013).

6. Gardiner, A. S., Twiss, J. L. & Perrone-Bizzozero, N. I. Competing interactions of RNA-binding proteins, Micrornas, and their targets control neuronal development and function. Biomolecules 5, 2903–2918 (2015).

7. Heraud-Farlow, J. E. et al. Staufen2 regulates neuronal target RNAs. Cell Rep 5, 1511–1518 (2013).

8. Lennox, A. L., Mao, H. & Silver, D. L. RNA on the brain: emerging layers of post- transcriptional regulation in cerebral cortex development. Wiley Interdiscip Rev Dev Biol 7, e290 (2018).

9. Naef, V. et al. The Stemness Gene Mex3A Is a Key Regulator of Neuroblast Proliferation During Neurogenesis. Front Cell Dev Biol 8, 549533 (2020).

10. Park, Y., Page, N., Salamon, I., Li, D. & Rasin, M. R. Making sense of mRNA landscapes: Translation control in neurodevelopment. Wiley Interdiscip Rev RNA 13, 1–22 (2022).

11. Weyn-Vanhentenryck, S. M. et al. Precise temporal regulation of alternative splicing during neural development. Nature Communications 2018 9:1 9, 1–17 (2018).

12. Zhou, Y., Dong, F. & Mao, Y. Control of CNS Functions by RNA-Binding Proteins in Neurological Diseases. Curr Pharmacol Rep 4, 301–313 (2018).

13. Schieweck, R., Ninkovic, J. & Kiebler, M. A. RNA-BINDING PROTEINS BALANCE BRAIN FUNCTION IN HEALTH AND DISEASE. Physiol Rev 101, 1309–1370 (2021).

14. Venables, J. P. et al. MBNL1 and RBFOX2 cooperate to establish a splicing programme involved in pluripotent stem cell differentiation. Nat Commun 4, 1–10 (2013).

15. Prashad, S. & Gopal, P. P. RNA-binding proteins in neurological development and disease. RNA Biol 18, 972–987 (2021).

16. Vessey, J. P. et al. An asymmetrically localized Staufen2-dependent RNA complex regulates maintenance of mammalian neural stem cells. Cell Stem Cell 11, 517–528 (2012).

17. Kusek, G. et al. Asymmetric segregation of the double-stranded RNA binding protein Staufen2 during mammalian neural stem cell divisions promotes lineage progression. Cell Stem Cell 11, 505–516 (2012).

18. Broadus, J., Fuerstenberg, S. & Doe, C. Q. Staufen-dependent localization of prospero mRNA contributes to neuroblast daughter-cell fate. Nature 391, 792–795 (1998).

19. Duchaîne, T. F. et al. Staufen2 isoforms localize to the somatodendritic domain of neurons and interact with different organelles. J Cell Sci 115, 3285–3295 (2002).

20. Zhong, W. & Chia, W. Neurogenesis and asymmetric cell division. Curr Opin Neurobiol **18**, 4–11 (2008).

21. Chowdhury, R. et al. STAU2 binds a complex RNA cargo that changes temporally with production of diverse intermediate progenitor cells during mouse corticogenesis. Development (Cambridge) 148, (2021).

22. Haubensak, W., Attardo, A., Denk, W. & Huttner, W. B. Neurons arise in the basal neuroepithelium of the early mammalian telencephalon: A major site of neurogenesis. Proc Natl Acad Sci U S A 101, 3196–3201 (2004).

23. Park, E., Gleghorn, M. L. & Maquat, L. E. Staufen2 functions in Staufen1-mediated mRNA decay by binding to itself and its paralog and promoting UPF1 helicase but not ATPase activity. Proc Natl Acad Sci U S A 110, 405–412 (2013).

24. Takahashi, K. & Yamanaka, S. Induction of Pluripotent Stem Cells from Mouse Embryonic and Adult Fibroblast Cultures by Defined Factors. Cell 126, 663–676 (2006).

25. Yan, Y. et al. Directed Differentiation of Dopaminergic Neuronal Subtypes from Human Embryonic Stem Cells. Stem Cells 23, 781–790 (2005).

26. Zeng, H. et al. Specification of Region-Specific Neurons Including Forebrain Glutamatergic Neurons from Human Induced Pluripotent Stem Cells. PLoS One 5, e11853 (2010).

27. Anderson, S. & Vanderhaeghen, P. Cortical neurogenesis from pluripotent stem cells: complexity emerging from simplicity. Curr Opin Neurobiol 27, 151–157 (2014).

28. Heber, S. et al. Staufen2-mediated RNA recognition and localization requires combinatorial action of multiple domains. Nature Communications 2019 10:1 10, 1–16 (2019).

29. Shi, Y., Kirwan, P. & Livesey, F. J. Directed differentiation of human pluripotent stem cells to cerebral cortex neurons and neural networks. Nat Protoc 7, 1836–1846 (2012).

30. Hao, Y. et al. Integrated analysis of multimodal single-cell data. Cell 184, 3573– 3587.e29 (2021).

31. Hao, Y. et al. Dictionary learning for integrative, multimodal and scalable single-cell analysis. Nat Biotechnol 42, 293–304 (2024).

32. Büttner, M., Ostner, J., Müller, C. L., Theis, F. J. & Schubert, B. scCODA is a Bayesian model for compositional single-cell data analysis. Nat Commun 12, 6876 (2021).

33. Finak, G. et al. MAST: a flexible statistical framework for assessing transcriptional changes and characterizing heterogeneity in single-cell RNA sequencing data. Genome Biol 16, 278 (2015).

34. Nguyen, H. C. T., Baik, B., Yoon, S., Park, T. & Nam, D. Benchmarking integration of single-cell differential expression. Nat Commun 14, 1570 (2023).

35. Subramanian, A., Kuehn, H., Gould, J., Tamayo, P. & Mesirov, J. P. GSEA-P: a desktop application for Gene Set Enrichment Analysis. Bioinformatics 23, 3251–3253 (2007).

36. Genaro-Mattos, T. C., Anderson, A., Allen, L. B., Korade, Z. & Mirnics, K. Cholesterol Biosynthesis and Uptake in Developing Neurons. ACS Chem Neurosci 10, 3671– 3681 (2019).

37. Zheng, X. et al. Metabolic reprogramming during neuronal differentiation from aerobic glycolysis to neuronal oxidative phosphorylation. Elife 5, (2016).

38. Rumpf, S., Sanal, N. & Marzano, M. Energy metabolic pathways in neuronal development and function. Oxford Open Neuroscience 2, 1–7 (2023).

39. Coolen, M., Katz, S. & Bally-Cuif, L. miR-9: A versatile regulator of neurogenesis. Front Cell Neurosci 7, 71003 (2013).

40. McGeary, S. E. et al. The biochemical basis of microRNA targeting efficacy. Science (1979) 366, eaav1741 (2019).

41. Brett, J. O., Renault, V. M., Rafalski, V. A., Webb, A. E. & Brunet, A. The microRNA cluster miR-106b∼25 regulates adult neural stem/progenitor cell proliferation and neuronal differentiation. Aging 3, 108–124 (2011).

42. de Sande, B. et al. A scalable SCENIC workflow for single-cell gene regulatory network analysis. Nat Protoc 15, 2247–2276 (2020).

43. Ramos, C., Fernández-Llebrez, P., Bach, A., Robert, B. & Soriano, E. Msx1 disruption leads to diencephalon defects and hydrocephalus. Developmental Dynamics 230, 446–460 (2004).

44. Ruzhynsky, V. A. et al. Cell Cycle Regulator E2F4 Is Essential for the Development of the Ventral Telencephalon. Journal of Neuroscience 27, 5926–5935 (2007).

45. Messmer, K., Shen, W. Bin, Remington, M. & Fishman, P. S. Induction of neural differentiation by the transcription factor NeuroD2. International Journal of Developmental Neuroscience 30, 105–112 (2012).

46. Moon, J. H. et al. Reprogramming of mouse fibroblasts into induced pluripotent stem cells with Nanog. Biochem Biophys Res Commun 431, 444–449 (2013).

47. Shen, T., Ji, F., Yuan, Z. & Jiao, J. CHD2 is Required for Embryonic Neurogenesis in the Developing Cerebral Cortex. Stem Cells 33, 1794–1806 (2015).

48. Angelis, N. et al. Loss of ARID3A perturbs intestinal epithelial proliferation– differentiation ratio and regeneration. Journal of Experimental Medicine 221, e20232279 (2024).

49. Shen, Z. et al. Enforcement of developmental lineage specificity by transcription factor Oct1. Elife 6, (2017).

50. Zhang, X. et al. Pax6 is a human neuroectoderm cell fate determinant. Cell Stem Cell 7, 90 (2010).

51. Ang, Y. S. et al. Disease Model of GATA4 Mutation Reveals Transcription Factor Cooperativity in Human Cardiogenesis. Cell 167, 1734–1749.e22 (2016).

52. Liu, J. C., Ferreira, C. G. & Yusufzai, T. Human CHD2 is a chromatin assembly ATPase regulated by its chromo- and DNA-binding domains. J Biol Chem 290, 25–34 (2015).

53. Carvill, G. L. et al. Targeted resequencing in epileptic encephalopathies identifies de novo mutations in CHD2 and SYNGAP1. Nat Genet 45, 825–830 (2013).

54. Folmes, C. D. L. et al. Somatic oxidative bioenergetics transitions into pluripotency- dependent glycolysis to facilitate nuclear reprogramming. Cell Metab 14, 264–271 (2011).

55. Folmes, C. D. L., Dzeja, P. P., Nelson, T. J. & Terzic, A. Metabolic plasticity in stem cell homeostasis and differentiation. Cell Stem Cell 11, 596–606 (2012).

56. Zhang, J., Nuebel, E., Daley, G. Q., Koehler, C. M. & Teitell, M. A. Metabolic regulation in pluripotent stem cells during reprogramming and self-renewal. Cell Stem Cell 11, 589–595 (2012).

57. Qi, C. et al. Hypoxia stimulates neural stem cell proliferation by increasing HIF"‘1Î± expression and activating Wnt/Î^2^-catenin signaling. Cell Mol Biol 63, 12–19 (2017).

58. Licausi, F. & Hartman, N. W. Role of mTOR Complexes in Neurogenesis. International Journal of Molecular Sciences 2018, Vol. 19, Page 1544 19, 1544 (2018).

59. Ho, J. J. D. et al. A network of RNA-binding proteins controls translation efficiency to activate anaerobic metabolism. Nature Communications 2020 11:1 11, 1–16 (2020).

60. Ebert, M. S. & Sharp, P. A. Roles for MicroRNAs in conferring robustness to biological processes. Cell 149, 515–524 (2012).

61. Delaloy, C. et al. MicroRNA-9 Coordinates Proliferation and Migration of Human Embryonic Stem Cell-Derived Neural Progenitors. Cell Stem Cell 6, 323–335 (2010).

62. Zhao, C., Sun, G., Li, S. & Shi, Y. A feedback regulatory loop involving microRNA-9 and nuclear receptor TLX in neural stem cell fate determination. Nat Struct Mol Biol 16, 365–371 (2009).

63. Tan, X. et al. The CREB-miR-9 Negative Feedback Minicircuitry Coordinates the Migration and Proliferation of Glioma Cells. PLoS One 7, e49570 (2012).

64. Jiang, T. et al. MicroRNA-218 regulates neuronal radial migration and morphogenesis by targeting Satb2 in developing neocortex. Biochem Biophys Res Commun 647, 9– 15 (2023).

65. Shibata, M., Nakao, H., Kiyonari, H., Abe, T. & Aizawa, S. MicroRNA-9 Regulates Neurogenesis in Mouse Telencephalon by Targeting Multiple Transcription Factors. Journal of Neuroscience 31, 3407–3422 (2011).

66. Velasco, M. X. et al. Antagonism between the RNA-binding protein Musashi1 and miR-137 and its potential impact on neurogenesis and glioblastoma development. RNA 27, 768–782 (2019).

67. Choudhury, N. R. et al. Tissue-specific control of brain-enriched miR-7 biogenesis. Genes Dev 27, 24–38 (2013).

68. Semba, Y. et al. Chd2 regulates chromatin for proper gene expression toward differentiation in mouse embryonic stem cells. Nucleic Acids Res 45, 8758–8772 (2017).

69. Rhee, C. et al. Arid3a is essential to execution of the first cell fate decision via direct embryonic and extraembryonic transcriptional regulation. Genes Dev 28, 2219–2232 (2014).

70. Gutiérrez-Franco, A. et al. Methanol fixation is the method of choice for droplet-based single-cell transcriptomics of neural cells. Communications Biology 2023 6:1 6, 1–12 (2023).

71. Macosko, E. Z. Z. et al. Highly parallel genome-wide expression profiling of individual cells using nanoliter droplets. Cell 161, 1202–1214 (2015).

72. Bio, D. Dolomite Bio Nadia Instrument Application Note for scRNA-seq Encapsulating single cells with barcoded beads on the Nadia Instrument.

73. Drop-seq tools: Java tools for analyzing Drop-seq data. https://github.com/broadinstitute/Drop-seq.

74. Zerbino, D. R. et al. Ensembl 2018. Nucleic Acids Res 46, D754–D761 (2018).

75. Picard Tools - By Broad Institute. https://broadinstitute.github.io/picard/.

76. Dobin, A. et al. STAR: ultrafast universal RNA-seq aligner. Bioinformatics 29, 15–21 (2013).

77. Hao, Y. et al. Integrated analysis of multimodal single-cell data. Cell 0, 3573–3587 (2021).

78. Luecken, M. D. & Theis, F. J. Current best practices in single-cell RNA-seq analysis: a tutorial. Mol Syst Biol 15, 8746 (2019).

79. McGinnis, C. S., Murrow, L. M. & Gartner, Z. J. DoubletFinder: Doublet Detection in Single-Cell RNA Sequencing Data Using Artificial Nearest Neighbors. Cell Syst 8, 329–337.e4 (2019).

80. McGinnis, C. S., Murrow, L. M. & Gartner, Z. J. DoubletFinder: Doublet Detection in Single-Cell RNA Sequencing Data Using Artificial Nearest Neighbors. Cell Syst 8, 329–337.e4 (2019).

81. Liberzon, A. et al. Molecular signatures database (MSigDB) 3.0. Bioinformatics 27, 1739–1740 (2011).

82. Liberzon, A. et al. The Molecular Signatures Database (MSigDB) hallmark gene set collection. Cell Syst 1, 417 (2015).

83. Subramanian, A. et al. Gene set enrichment analysis: A knowledge-based approach for interpreting genome-wide expression profiles. Proc Natl Acad Sci U S A 102, 15545–15550 (2005).

84. Sayols, S. rrvgo: a Bioconductor package for interpreting lists of Gene Ontology terms. MicroPubl Biol 2023, (2023).

85. Seal, R. L. et al. Genenames.org: the HGNC resources in 2023. Nucleic Acids Res 51, D1003–D1009 (2023).

86. Harrison, P. W. et al. Ensembl 2024. Nucleic Acids Res 52, D891–D899 (2024).

87. Walsh, R. M. et al. Generation of human cerebral organoids with a structured outer subventricular zone. Cell Rep 43, 114031 (2024).

