## Supplementary Material for "Staufen2 modulates the temporal dynamics of human neurogenesis *in vitro*"

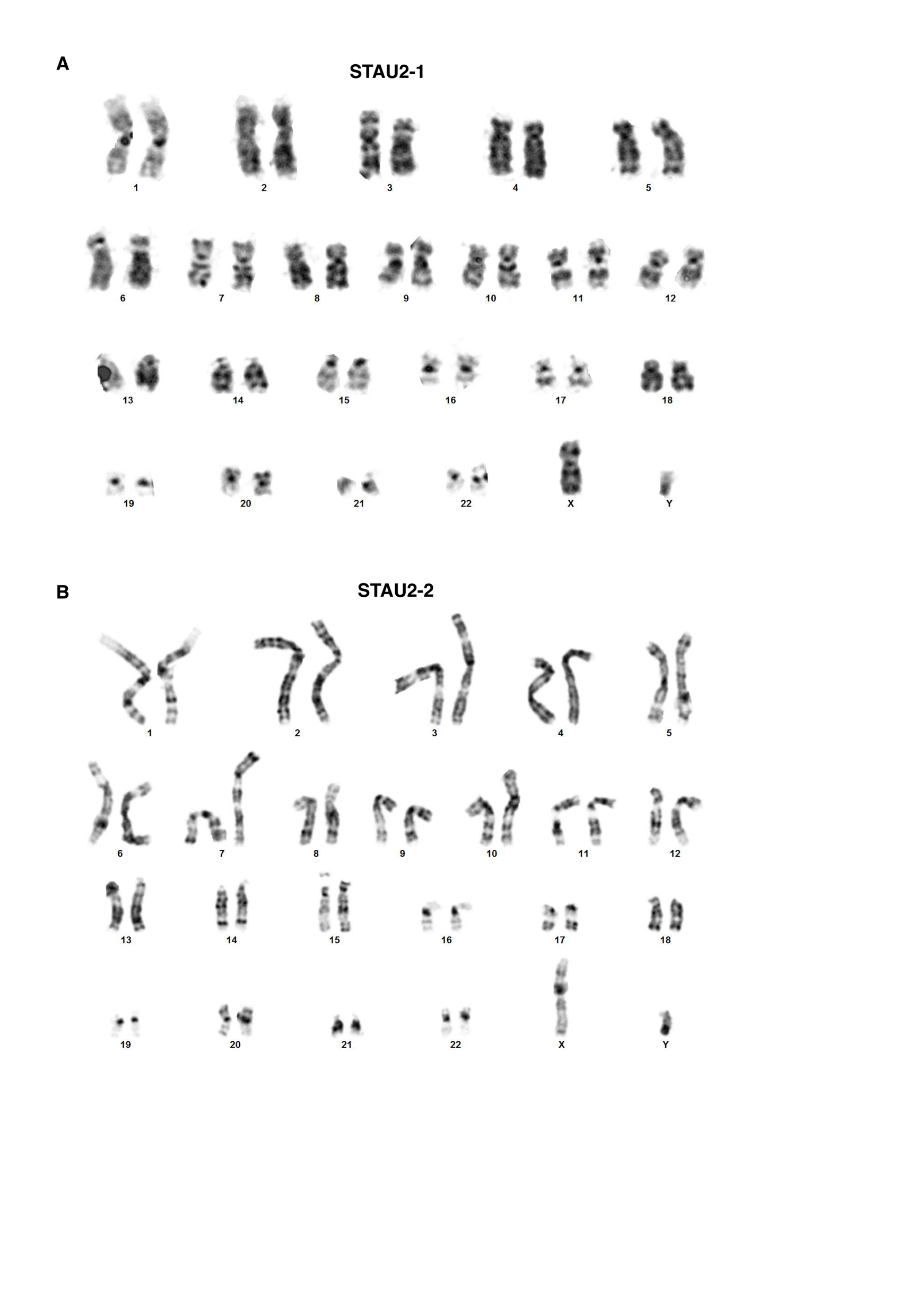

**Figure S1. Karyotypes of STAU2 KO cell lines. A, B.** Karyotyping of STAU2-1 (**A**) and STAU2-2 (**B**) reveals normal chromosomal integrity.

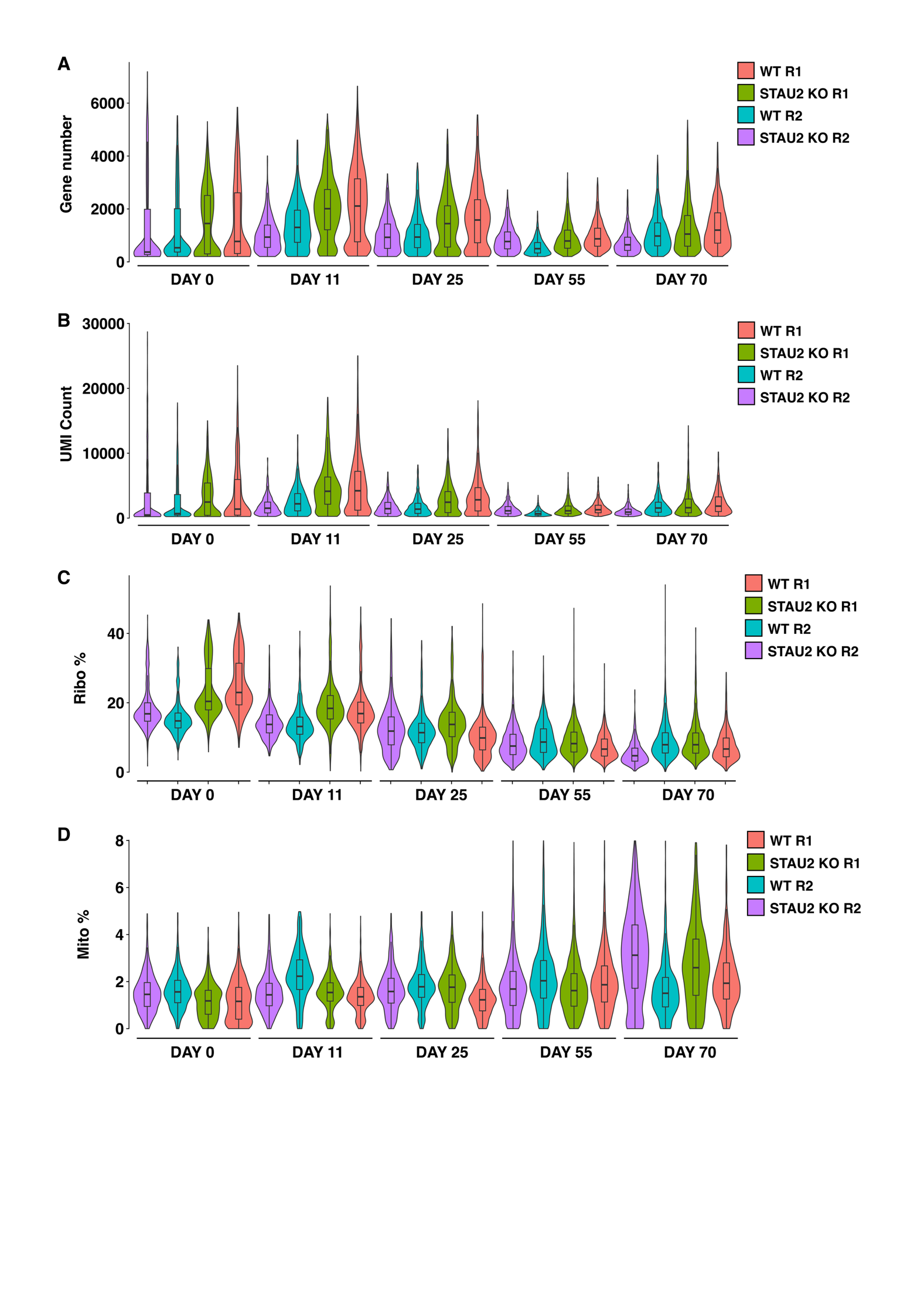
**Figure S2. Quality control plots of the single-cell data.** Violin plots depicting the distribution of genes per cell **(A)**, UMIs per cell **(B)**, the % of mitochondrial UMIs **(C)** and the % of Ribosomal UMIs **(D)** across all samples analyzed. Violin plots are colored by the sample they come as purple (Replicate 1 WT), cyan (Replicate 1 KO), olive green (Replicate 2 WT) and salmon (Replicate 2 KO)

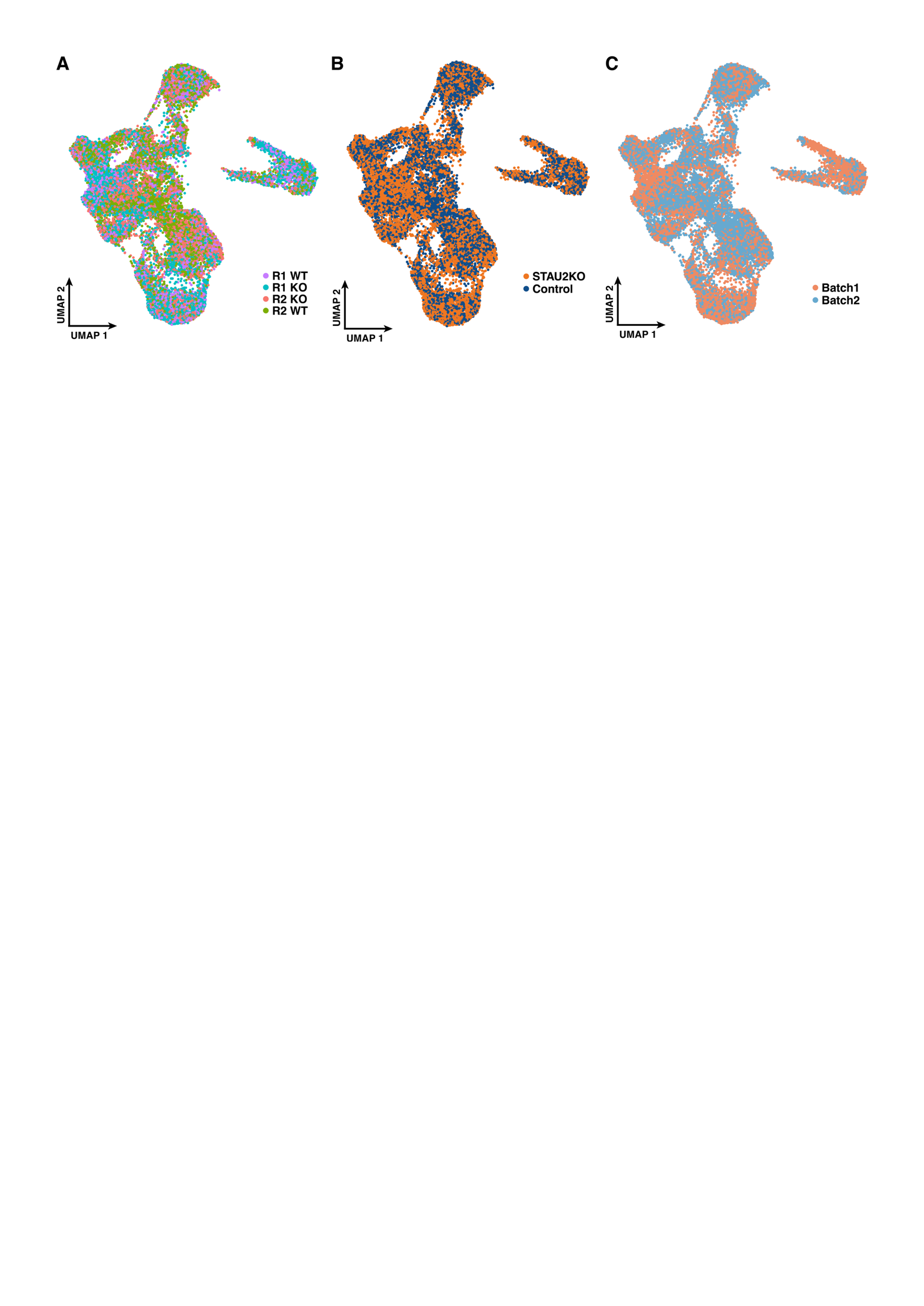

**Figure S3**. **Results of the integration of single-cell datasets using Canonical Correlation Analysis**. UMAP plots of the integrated dataset are colored by **(A)** sample (R1 WT, R1 KO, R2 WT, R2 KO), **(B)** condition (STAU2 KO vs. Control), and **(C)** batch (Batch 1 vs. Batch 2). Cell distribution is largely consistent across samples and conditions, indicating successful integration. While we do see minimal batch effects, it is important to not over-correct at the risk of losing biological variation.

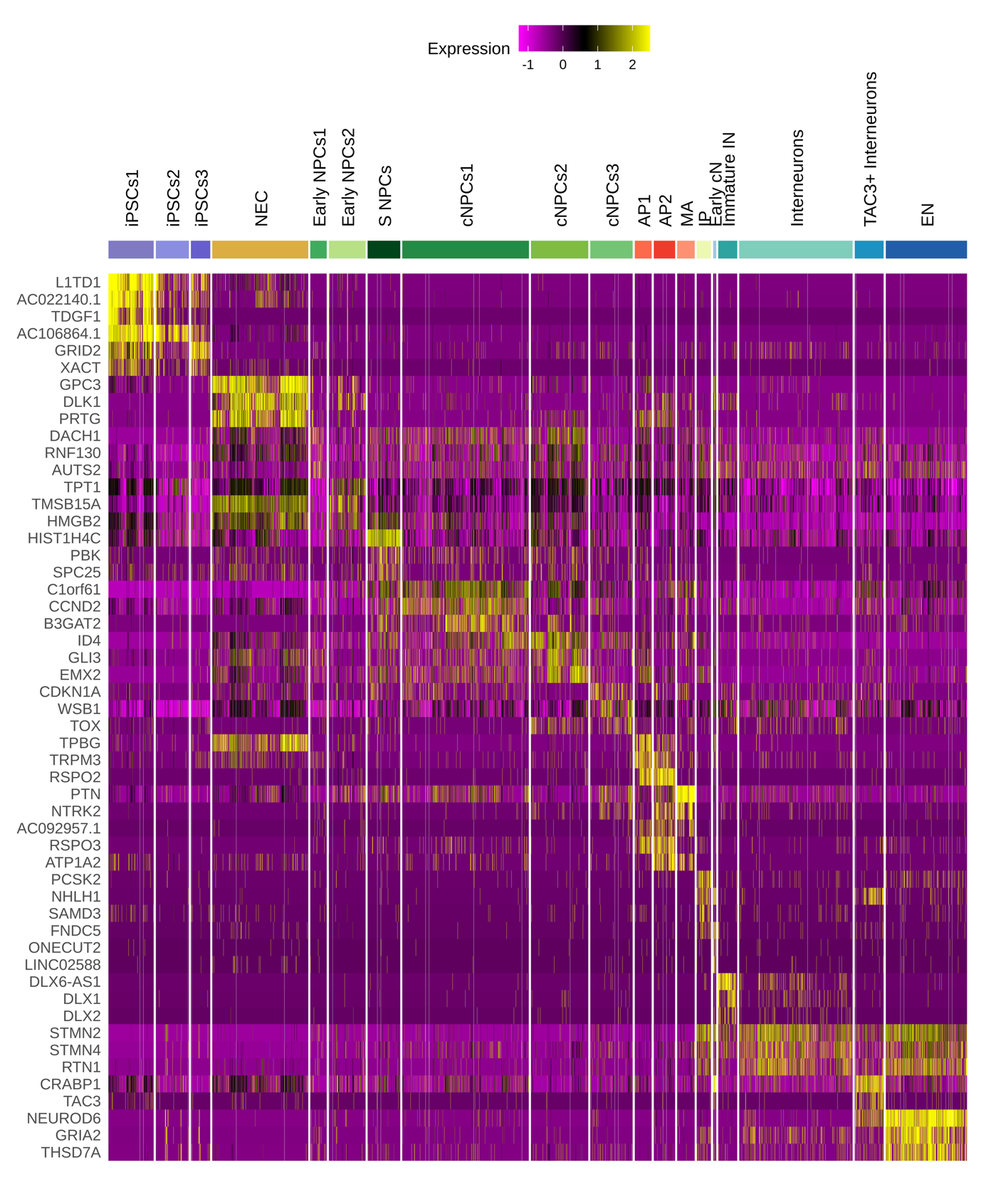

**Figure S4. Additional markers for each cluster identified in the scRNA-seq data.**  Heatmap displaying the top three marker genes for each cluster identified in the scRNA-seq dataset. Gene expression values are row-normalized to allow comparisons across clusters. Yellow indicates higher expression, and purple indicates lower expression. Clusters are annotated at the top and grouped by major cell types.

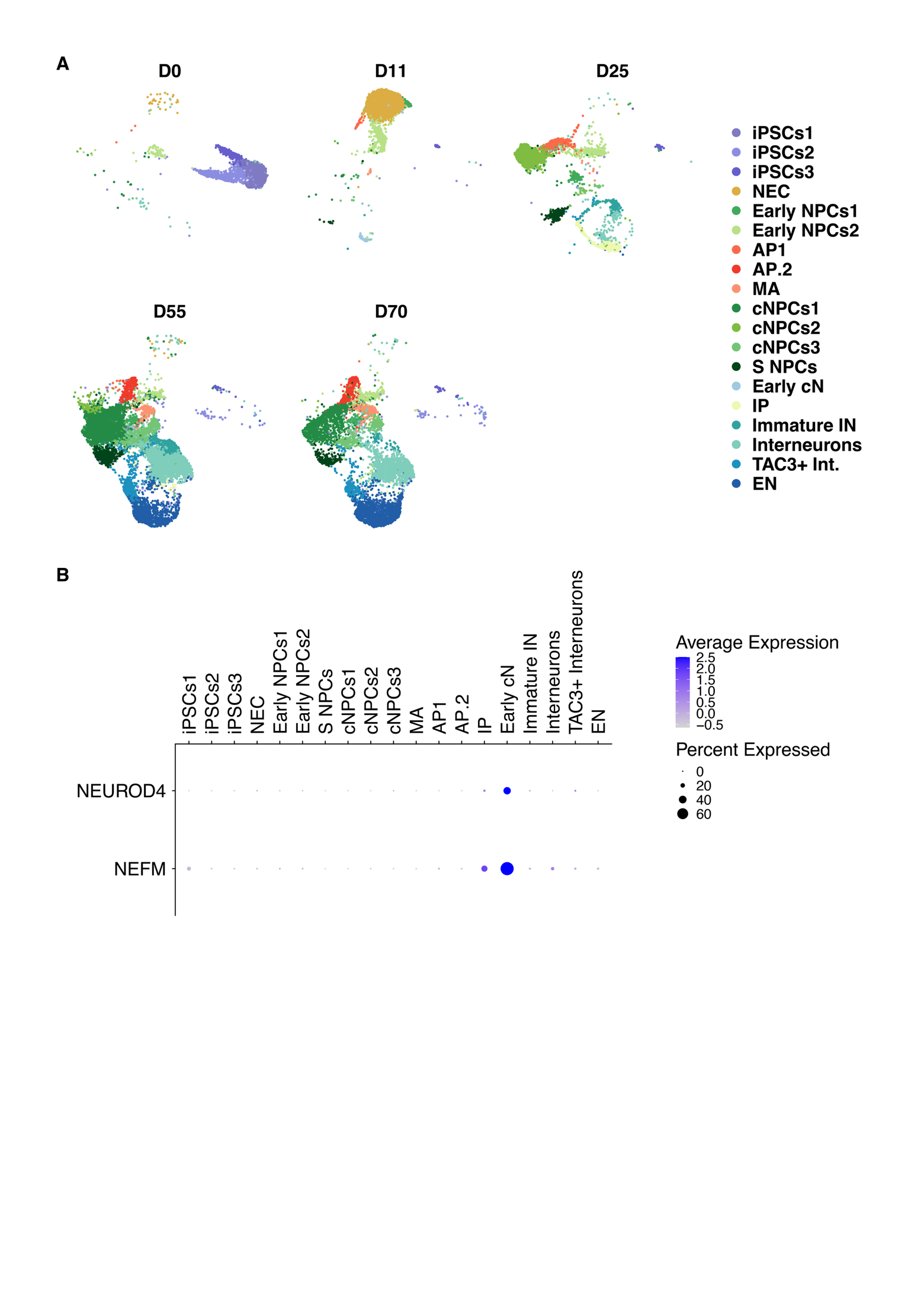

**Figure S5. Identified cell populations at different time points. A.** UMAP plots showing the cell clusters split by time point. **B.** Dot plot showing the expression of cortical marker genes NEUROD4 and NEFM across clusters. Dot size represents the percentage of cells expressing the gene in each cluster, while color intensity reflects the average expression level. Strong expression in the Early.Cortical.Neurons (ECN) cluster suggests a cortical identity for these cells.

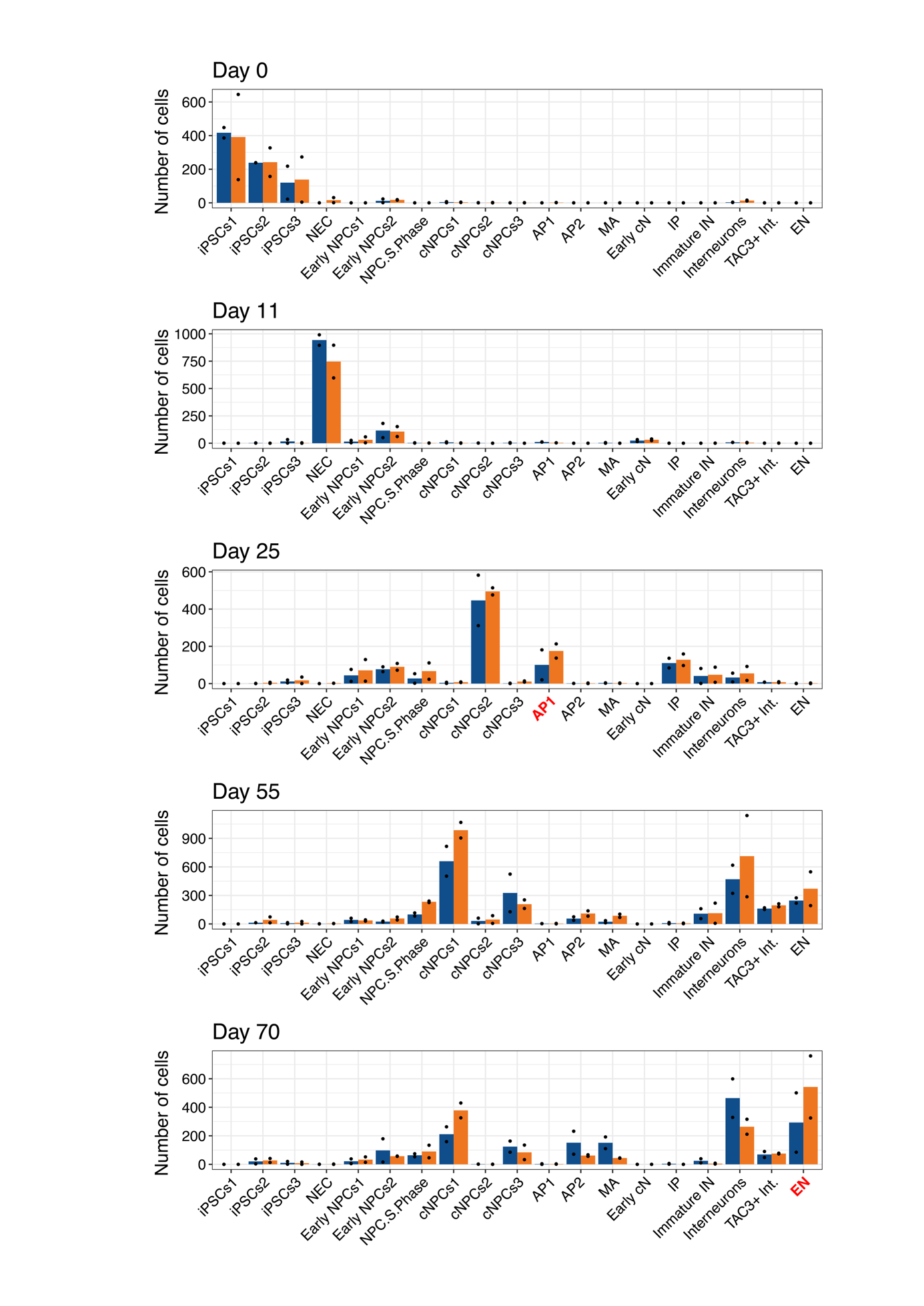

**Figure S6. Cell composition analysis using scCODA.** Barplot showing the proportion of each cell type in each time point. Bars represent the average of the two replicates and black dots the values of each of the samples. Cell clusters with significant changes in cell proportion according to scCODA are highlighted in red.

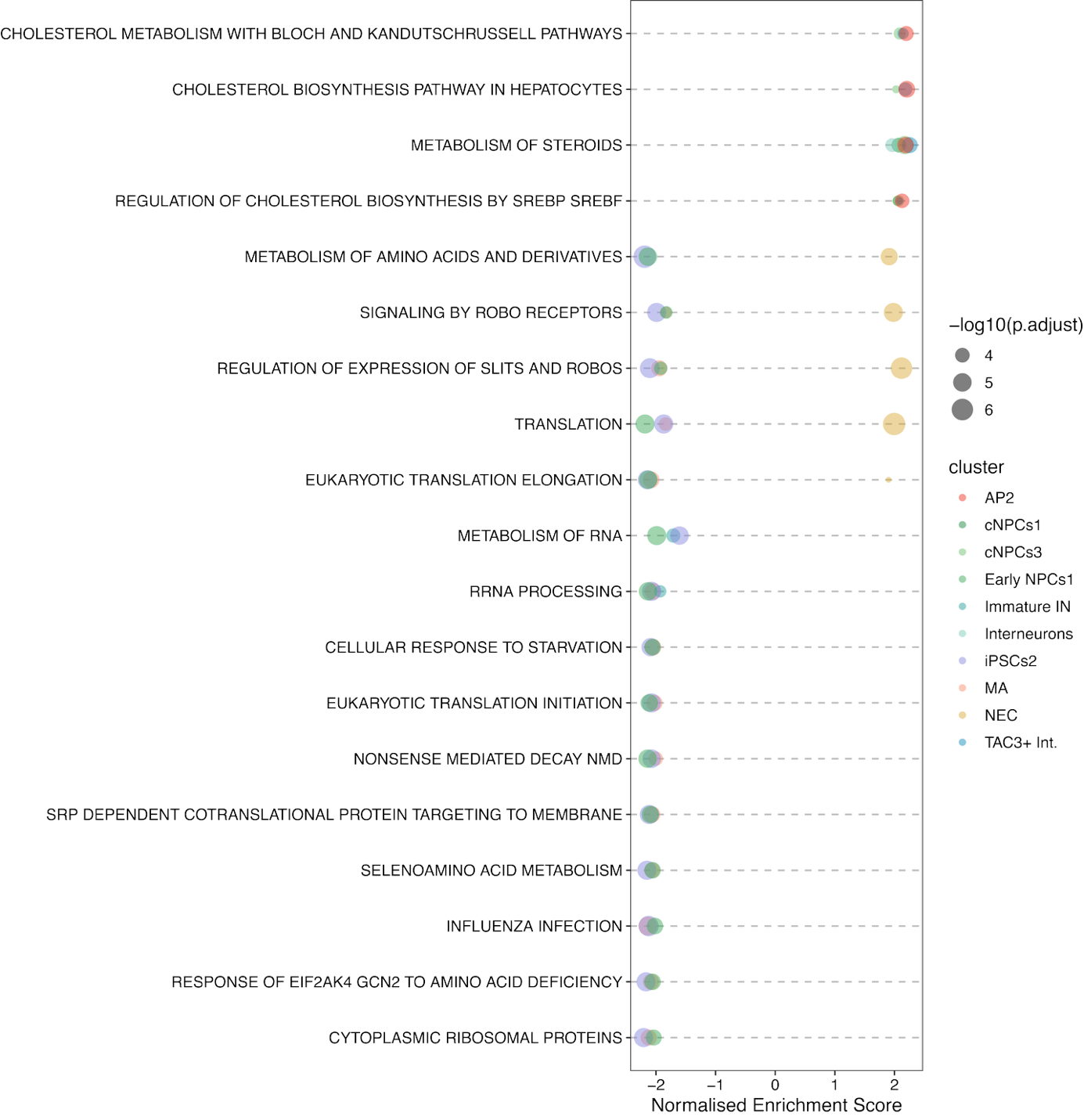

**Figure S7. Global impact of STAU2 across cell populations.** Bubble plot depicting pathways disregulated by STAU2 across multiple clusters (FDR < 0.001). Only pathways disregulated across 3 or more cell types are shown.

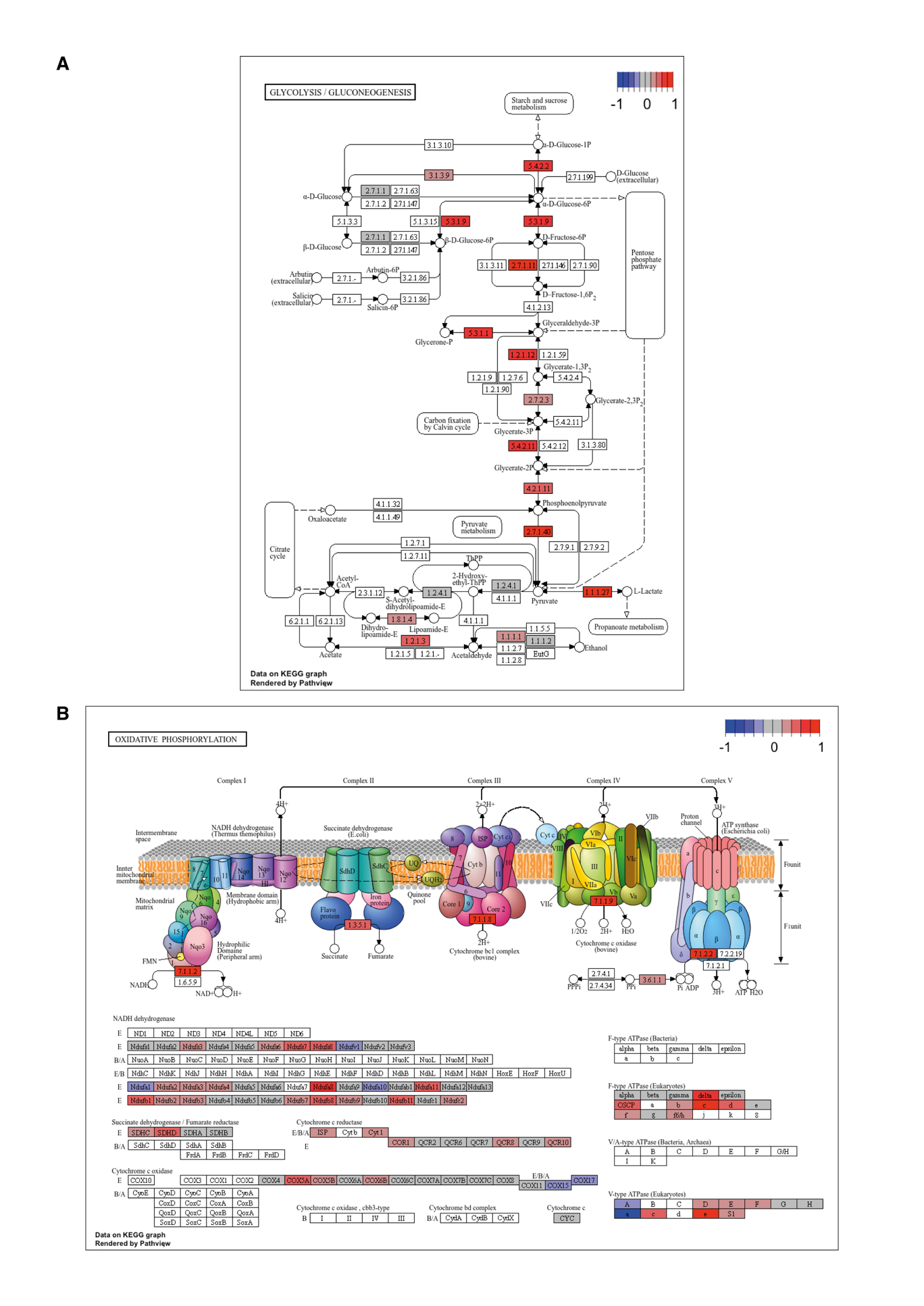

**Figure S8. Graphical representation of the effects of STAU2 on metabolic pathways in neuroepithelial cells.** Representation of the glycolysis/gluconeogenesis **(A)** and the oxidative phosphorylation **(B)** pathways according to KEGG. Changes in expression of specific enzymes in the pathways in STAU2 KO samples compared to WT is shown in colors to represent downregulated (blue) and upregulated (red) enzymes.

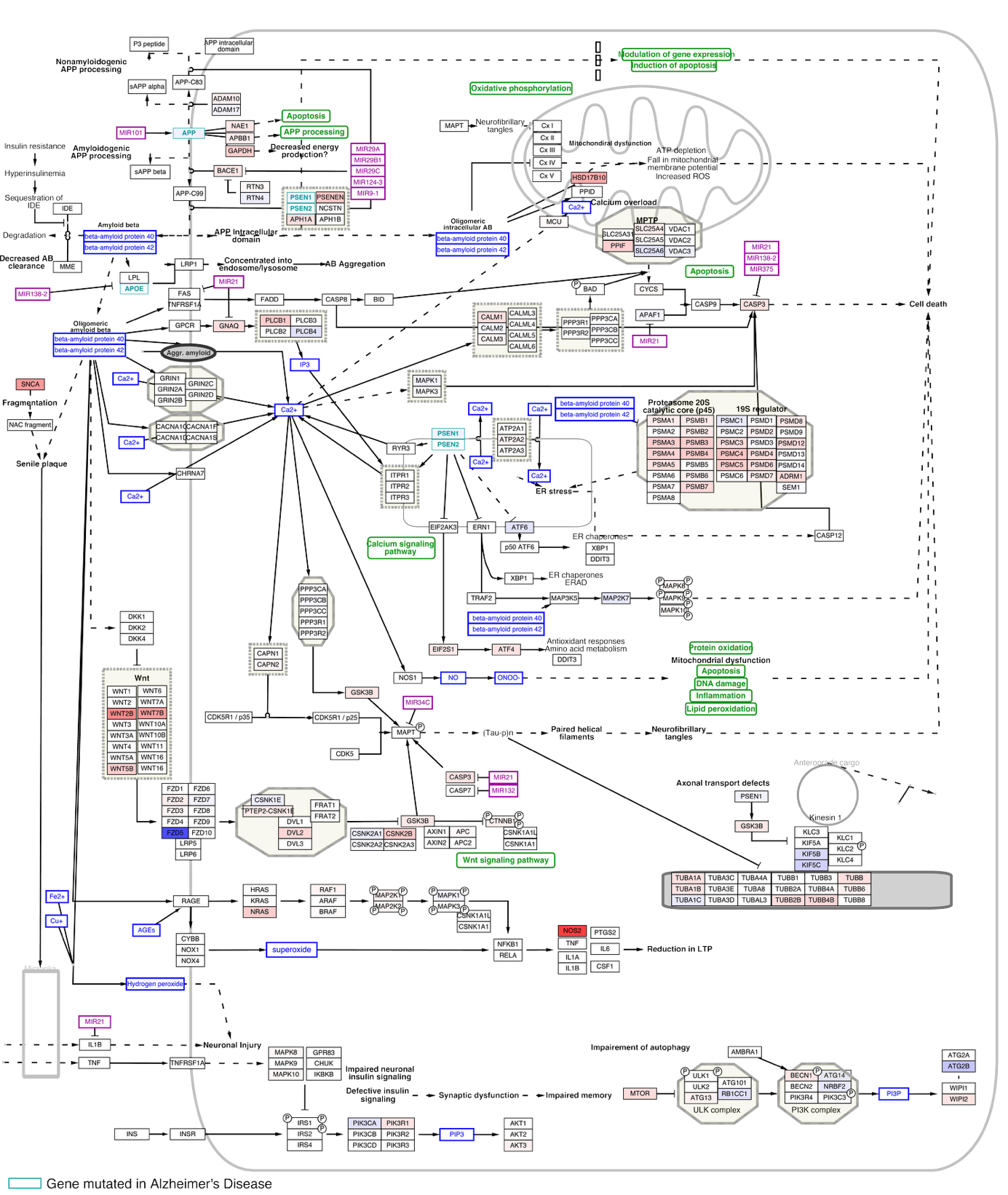

**Figure S9. Graphical representation of the effects of STAU2 on the pathway *Alzheimer’s Disease and miRNA effects* in neuroepithelial cells.** Changes in expression of specific enzymes in the pathways in STAU2 KO samples compared to WT is shown in colors to represent downregulated (blue) and upregulated (red) enzymes. Differentially regulated miRNAs in Alzheimer’s Disease are shown in purple.

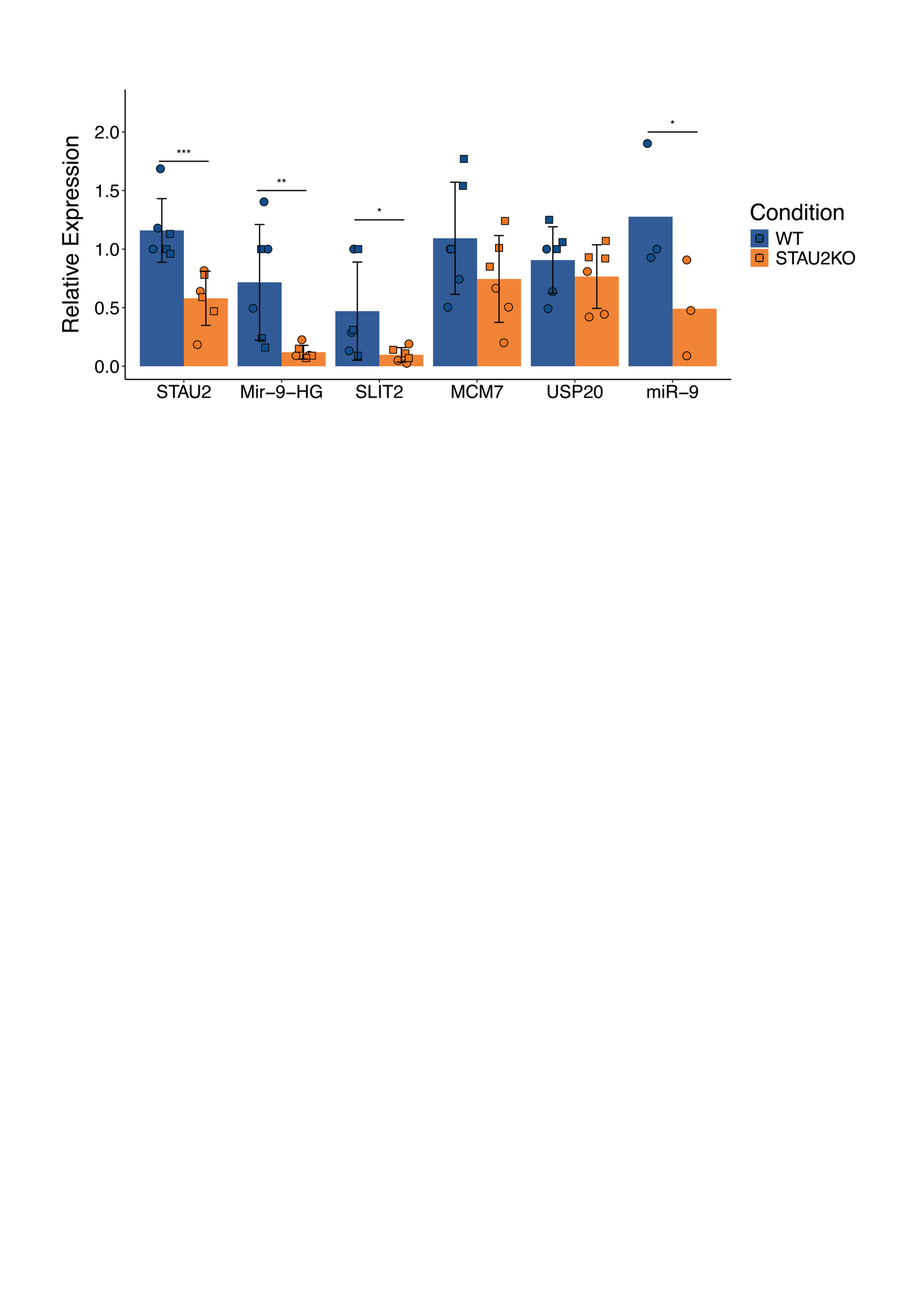

**Figure S10. qRT-PCR validation of miRNA host genes and mature miRNAs.** RT-PCR validation of selected miRNA host genes (MIR9HG-1, SLIT2, MCM7) in two independent STAU2 KO lines (denoted as circles and squares on the graph, respectively). TaqMan qPCR data for miR-9 were normalized using housekeeping RPL21 and RNU58B Ct values, whereas RT-qPCR data for long transcripts were normalized to GAPDH and HPRT housekeeping control genes. Downregulation of *MIR9HG-1* is accompanied by downregulation of its respective mature miR-9. Data are presented as mean ± SD; ** p-value < 0.01; *** p-value < 0.001, (Non-parametric Two-way ANOVA test, *n* = 3 biological replicates per genotype).

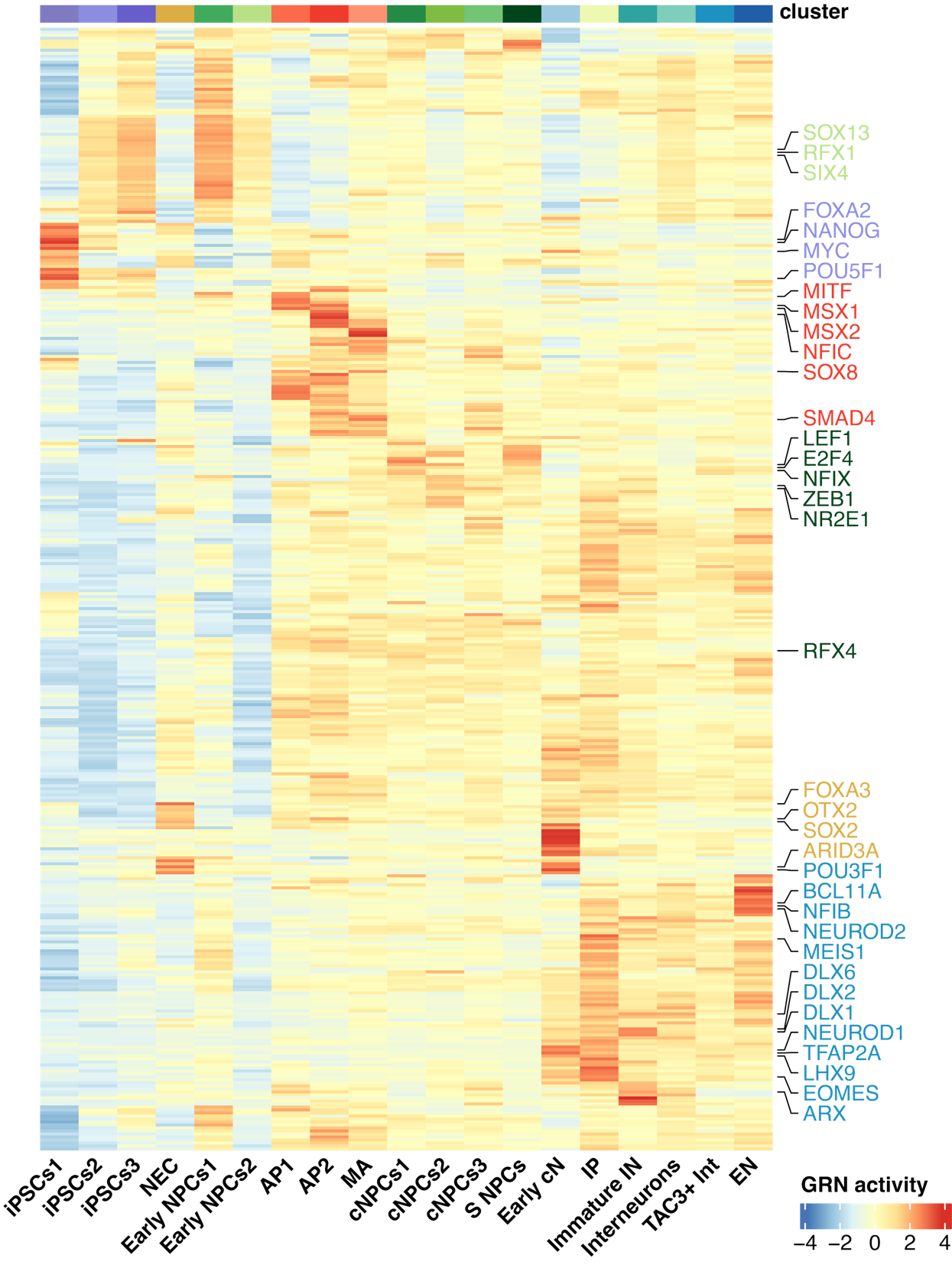

**Figure S11. Cell type specific transcriptional regulatory landscapes.** Heatmap showing the activity of gene regulatory networks (GRNs) inferred by SCENIC across the different cell clusters. Each column represents a cluster, and each row corresponds to a GRN, represented by its associated transcription factor. Transcription factors with enriched activity in specific cell types are highlighted: iPSCs (purple), astrocytes (red), progenitor cells (green), neuroepithelial cells (yellow), and neurons (blue).

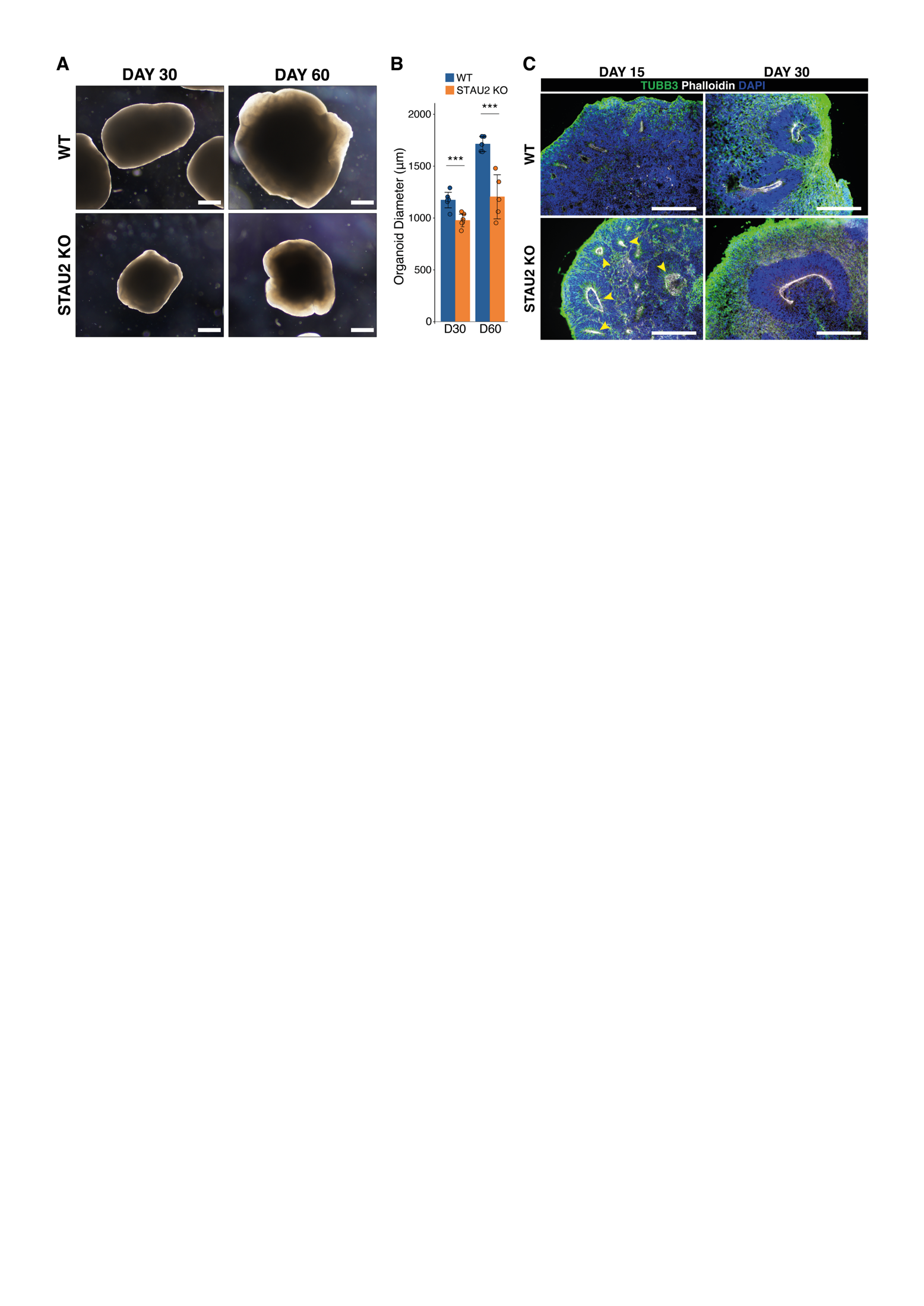

**Figure S12. STAU2-2 organoids show neurogenesis defects. A.** Images of STAU2-2 and WT2 organoids at 30 (D30) and 60 (D60) days**. B.** Quantification of organoid diameter at D30 and D60 reveals a significant reduction in STAU2-2 KO organoid size compared to the isogenic control line (WT). (Non-parametric two-way ANOVA test, p-value < 0.001; ***). **C.** Immunofluorescence analysis of cortical organoids stained for TBB3 (green, neuronal marker), DAPI (blue, nuclei), and Phalloidin (white, labeling actin-rich neural rosettes). At D30, STAU2 KO organoids show increased neuronal differentiation and larger, more prominent neural rosettes compared to WT. By D60, the difference in neuronal density is reduced.

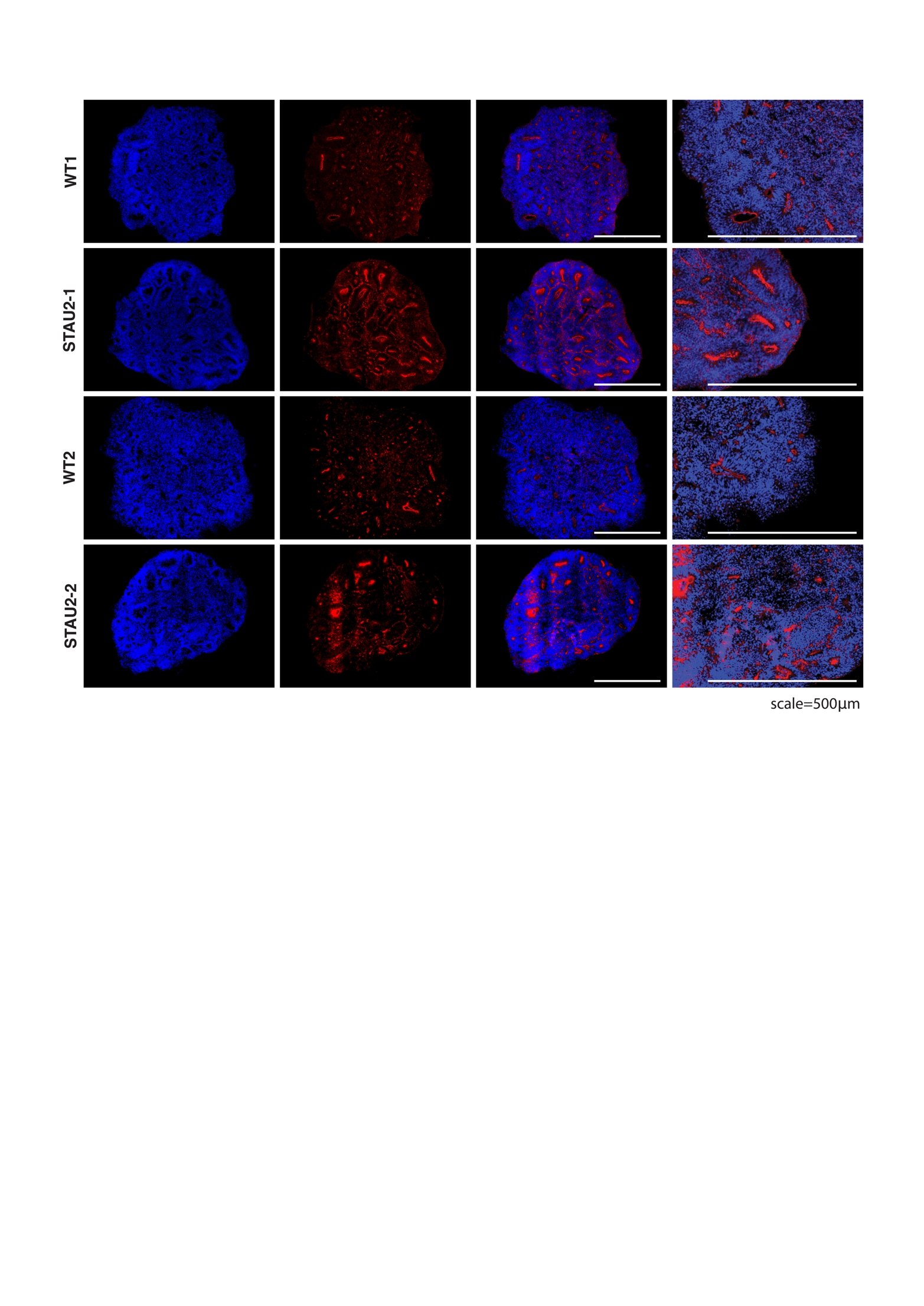

**Figure S13. Immunofluorescence analysis of ZO-1** **neuroepithelial marker in D15 cortical organoids.** Immunofluorescence staining of organoid sections from two wild type (WT1, WT2) and two STAU2 KO (STAU2-1, STAU2-2) lines. Blue indicates nuclear staining (DAPI), and red represents immunostaining for ZO-1, a tight-junction marker expressed in neuroepithelial cells. Columns show full organoid sections and zoomed-in views highlighting structural organization. Scale bars: 500 μm.

**Supplementary References**

83. Darmanis, S. et al. A survey of human brain transcriptome diversity at the single cell level. Proc Natl Acad Sci U S A 112, 7285–7290 (2015).
84. Ito, D. et al. Microglia-specific localisation of a novel calcium binding protein, Iba1. Molecular Brain Research 57, 1–9 (1998).
85. Lake, B. B. et al. Integrative single-cell analysis of transcriptional and epigenetic states in the human adult brain. Nat Biotechnol 36, 70–80 (2018).
86. Ayhan, F. et al. Resolving cellular and molecular diversity along the hippocampal anterior-to-posterior axis in humans. Neuron 109, 2091-2105.e6 (2021).
87. Su, Y. et al. A single-cell transcriptome atlas of glial diversity in the human hippocampus across the postnatal lifespan. Cell Stem Cell 29, 1594-1610.e8 (2022).
88. Tasic, B. et al. Shared and distinct transcriptomic cell types across neocortical areas. Nature 563, 72–78 (2018).
89. Seeker, L. A. et al. Brain matters: unveiling the distinct contributions of region, age, and sex to glia diversity and CNS function. Acta Neuropathol Commun 11, 84 (2023).
90. Tran, M. N. et al. Single-nucleus transcriptome analysis reveals cell-type-specific molecular signatures across reward circuitry in the human brain. Neuron 109, 3088-3103.e5 (2021).
91. Beletskiy, A. et al. Detailed Analysis of Dorsal-Ventral Gradients of Gene Expression in the Hippocampus of Adult Rats. Int J Mol Sci 23, 9948 (2022).
92. Garma, L. D. et al. Interneuron diversity in the human dorsal striatum. (2023).
93. Zhong, S. et al. A single-cell RNA-seq survey of the developmental landscape of the human prefrontal cortex. Nature 555, 524–528 (2018).
94. Franjic, D. et al. Transcriptomic taxonomy and neurogenic trajectories of adult human, macaque, and pig hippocampal and entorhinal cells. Neuron 110, 452-469.e14 (2022).
95. de Ceglia, R. et al. Specialized astrocytes mediate glutamatergic gliotransmission in the CNS. Nature 2023 622:7981 622, 120–129 (2023).
96. Hodge, R. D. et al. Conserved cell types with divergent features in human versus mouse cortex. Nature 573, 61–68 (2019).
97. Lake, B. B. et al. Neuronal subtypes and diversity revealed by single-nucleus RNA sequencing of the human brain. Science (1979) 352, 1586–1590 (2016).
98. Zhou, Y. et al. Molecular landscapes of human hippocampal immature neurons across lifespan. Nature 607, 527–533 (2022).
99. Shin, J. et al. Single-Cell RNA-Seq with Waterfall Reveals Molecular Cascades underlying Adult Neurogenesis. Cell Stem Cell 17, 360–372 (2015).
100. Anthony, T. E., Klein, C., Fishell, G. & Heintz, N. Radial Glia Serve as Neuronal Progenitors in All Regions of the Central Nervous System. Neuron 41, 881–890 (2004).
101. Pollen, A. A. et al. Molecular Identity of Human Outer Radial Glia during Cortical Development. Cell 163, 55–67 (2015).
102. Habib, N. et al. Div-Seq: Single-nucleus RNA-Seq reveals dynamics of rare adult newborn neurons. Science (1979) 353, 925–928 (2016).
103. Cao, J. et al. The single-cell transcriptional landscape of mammalian organogenesis. Nature 566, 496–502 (2019).
104. Chamling, X. et al. Single-cell transcriptomic reveals molecular diversity and developmental heterogeneity of human stem cell-derived oligodendrocyte lineage cells. Nature Communications 2021 12:1 12, 1–20 (2021).
105. Zeisel, A. et al. Molecular Architecture of the Mouse Nervous System. Cell 174, 999-1014.e22 (2018).
